# A transcriptional cell atlas identifies the decline in the AT2 niche in aged human lungs

**DOI:** 10.1101/2023.06.16.545378

**Authors:** Xue Liu, Xuexi Zhang, ChangFu Yao, Jiurong Liang, Paul W. Noble, Dianhua Jiang

## Abstract

Aging poses a global public health challenge, associated with molecular and physiological changes in the lungs. It increases susceptibility to acute and chronic lung diseases, yet the underlying molecular and cellular drivers in aged populations are not fully appreciated. To systematically profile the genetic changes associated with age, we present a single-cell transcriptional atlas comprising nearly half a million cells from the healthy lungs of human subjects spanning various ages, sexes, and smoking statuses. Most annotated cell lineages in aged lungs exhibit dysregulated genetic programs. Specifically, the aged alveolar epithelial cells, including both alveolar type II (AT2) and type I (AT1) cells, demonstrate loss of epithelial identities, heightened inflammaging characterized by increased expression of AP-1 transcription factor and chemokine genes, and significantly increased cellular senescence. Furthermore, the aged mesenchymal cells display a remarkable decrease in Collagen and Elastin transcription. The decline of the AT2 niche is further exacerbated by a weakened endothelial cell phenotype and a dysregulated genetic program in macrophages. These findings highlight the dysregulation observed in both AT2 stem cells and their supportive niche cells, potentially contributing to the increased susceptibility of aged populations to lung diseases.

## INTRODUCTION

In the past century, human life expectancy has nearly doubled globally with an increase more than it did in all previous millennia combined. By 2019 over 600 million people are 65 years old or older and the number will likely triple by 2050 ^1^. Increases in human age necessitate new scientific understanding of how to both extend and enhance health over the course of human longer lives ^2^. Respiratory system, representing a unique interface with the outside environment, is one of the major organs that are reported to be most affected by human aging ^3^. Several lung diseases have aging as a major risk factor, including Chronic obstructive pulmonary disease (COPD), interstitial fibrotic lung disease (ILD), lung cancer and inflammatory lung diseases such as pneumonia and COVID-19 ^3,4^. However, it is not fully appreciated that what the mechanistic, molecular, and genetic drivers of lung aging are and how aging increases the susceptibility to these acute and chronic lung diseases.

Alveoli, the essential units for lung gas exchange, consist of epithelial and mesenchymal populations. Alveolar type II epithelial cells (AT2) have been well accepted as progenitor cells that maintain alveolar homeostasis and repair damaged epithelium after injury ^5^. Endothelial, immune, and mesenchymal components participate in the alveolar niche ^6–8^. However, AT2s exhibit reduced self-renewal and differentiation capacity with aging (PMID: 32591591). The aging of lung mesenchymal cells, vital for supporting AT2 renewal (PMID: 28886382), further impairs AT2 cell function (PMID: 34528872). However, a comprehensive dissection of the AT2 niche during lung aging using a large dataset is lacking.

Recent studies have been greatly benefited from the advancements of next-generation sequencing technologies, especially single-cell RNA sequencing (scRNA-seq), which has profiled healthy and disease human lungs ^9–13^. Specifically, the prevalence of the progressive lung diseases including idiopathic pulmonary fibrosis (IPF), COPD, COVID-19, and lung cancers increases with age. The hallmarks of these diseases, such as metabolic defects, genomic instability, telomere attritions, cellular senescence, genomic and epigenetic alterations, overlap with those that occur during aging and may act as sensitizers to these diseases ^14–24^. Although the single-cell transcriptomic atlas on these diseases have generate a huge amount of information on molecular and cellular profiles and disease specific alterations in the lungs, limited studies have been focused on aging of human lungs ^25,26^, and a longitudinal transcriptomic profiling of human lungs at single cell level across all adult age ranges is not available.

To fill the gap, here we generated a single-cell transcriptomic atlas of 491,187 human lung cells from 14 datasets with 92 adult healthy subjects of different ages, sexes, and smoking statuses. The comprehensive analyses elucidated the changes of transcriptional programs in all cell types in the lung during aging. Strikingly we found that a significant decline of the AT2 niche, characterized by the aberrantly inflammaging profile of AT2 themselves, and a decay of aged stromal cells in supporting alveolar epithelium. Dysregulated alveolar niches were further presented by impaired vascular and lymphatic endothelium and dysregulated transcriptional programs in both myeloid and lymphoid systems in aged lungs. Moreover, cell-cell communication analysis suggests a strong pro-inflammatory niche between AT2 cells and macrophages in aged subjects. Thus, these detailed observations highlight significant dysregulation in the aging of human lungs, shedding light on the mechanisms behind the decline in lung capacity, resilience, and increased susceptibility to diseases in older populations. The human lung atlas presented here serves as a valuable resource, resembling a pathogenetic-like library of aging lungs, and targeting specific molecular or cellular factors could potentially offer novel insights for intervening in age-related lung diseases.

## RESULTS

### Generation of the single cell transcriptional atlas on healthy human lungs

To generate a single-cell transcriptomic map on healthy human lungs and study the mechanisms underlying human lung aging, we utilized a good amount of single cell multiomics databases and extracted the data on healthy human lungs from 14 published databases ^10,12,14,24,27–35^ containing 94 healthy subjects and 491,187 cells (Figure 1A and Table S1). The donors were at wide age ranges (Figure S1A), both sexes, smoking histories, and different ethnic races, although most were Caucasians (Figure S1B).

**Figure 1.**
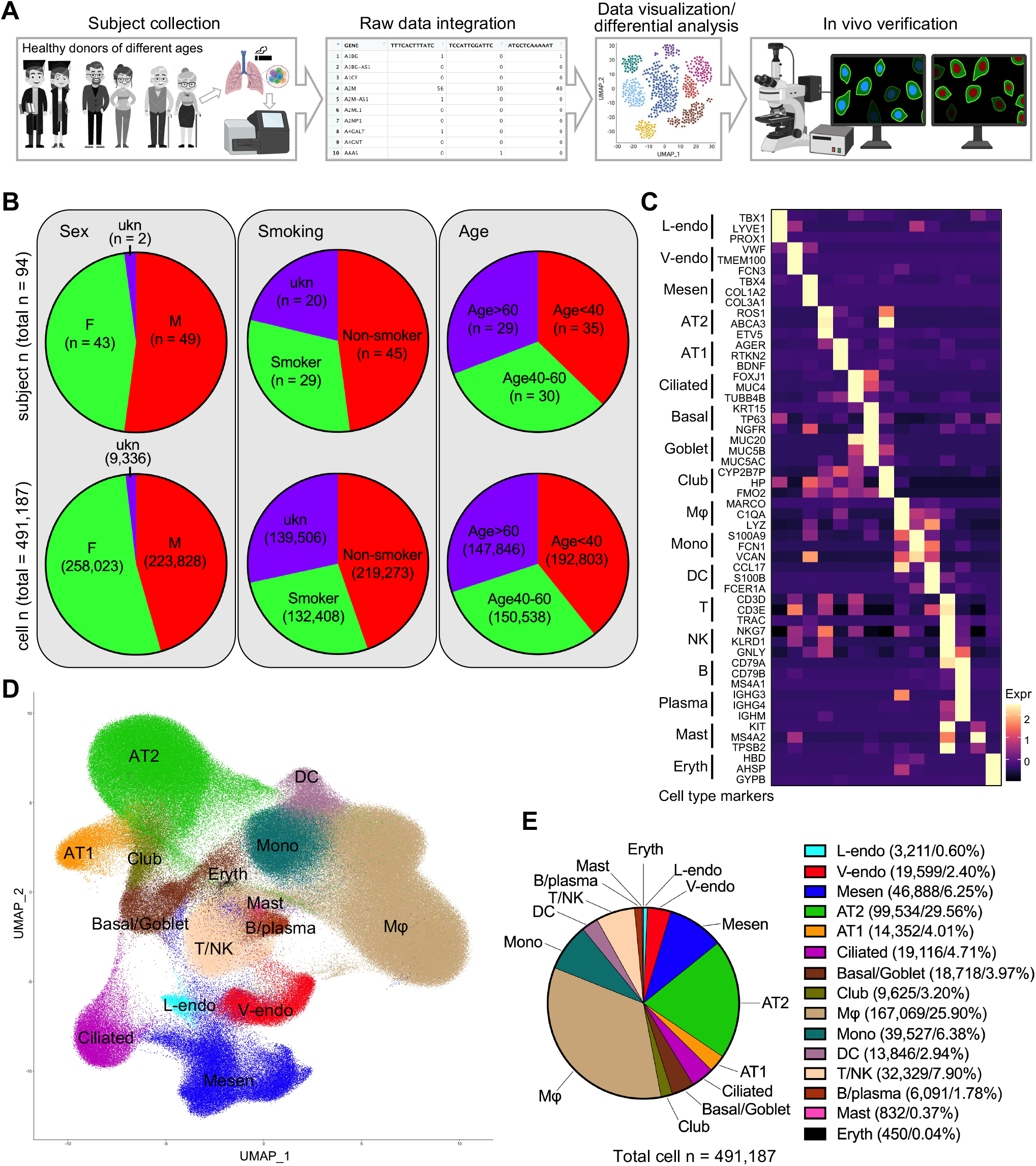
Integration of scRNA-seq data on human lungs from healthy subjects. **(A)** Schematic overview of the data collection, integration, differential analysis, and histological verifications of the healthy human lungs. The donors were all adults from different sex, age, smoking history. **(B)** The subjects and the cells were divided regarding to donor sex, smoking history, and age stages. For group by ages, the subjects were evenly divided into three group: age < 40, age 40-60, and age > 60. Pie charts were used to illustrate the corresponding subject/cell numbers and proportions. ukn, unknown. **(C)** The average transcriptions of the canonical cell type marker genes (rows) in cell types (columns) were determined and visualized by heatmap in the integrated data. **(D)** The distribution of the cells in the integrated data was visualized by UMAP and major cell types were identification based on the expression of the cell type marker genes in (**C**) and (S2**A**). **(E)** Cell number of each major cell type in the the integrated data was determined and the proportion of each cell type was quantified and visualized by pie chart. F, female, M, male, ukn, unknown. L/V-endo, lymphatic/vascular endothelial cells, Mesen, mesenchymal cells, AT1/2, alveolar type I/II epithelial cells, Mφ, macrophages, Mono, monocytes, DC, dendritic cells, Eryth, erythrocytes.

To study the lung transcriptional profiles of different conditions, the subjects and the cells were divided by the donor identities including sex (female, male, and unknown sex donors), smoking history (smoker, non-smoker, and smoking-history unknown donors) and age range (donors of age < 40 yr, age 40-60 yr, and age > 60 yr) (Figure 1B). The cells were then integrated, and cell distributions were visualized by Uniform Manifold Approximation and Projections (UMAPs). The comparable sequencing depth of different datasets (Figure S1C) were confirmed although the transcription of some genes in GSE135893 were far beyond the scale of other datasets (Figure S1D), so the following comparative analyses were done with this dataset separated unless mentioned. Well-performed batch corrections (Figures S1E-I) were confirmed, and the cells were then clustered (Figure S1J) in the integrated data. We then cataloged the cells into 15 major cell types annotated with the transcriptions of the canonical marker genes (Figures 1C and S2A), composed of 5 distinct lineages including endothelial (lymphatic/vascular endothelial cells), mesenchymal, epithelial (AT2, AT1, ciliated, basal/goblet, and club cells), myeloid (macrophages, monocytes, dendritic cells, mast cells, and erythrocytes), and lymphoid (T/NK and B/plasma cells) lineages (Figures 1D-E). Unique gene transcriptional patterns verified the annotation of the cell types (Figure S2B).

### Accelerated cellular senescence and loss of epithelial identities in aged AT2s

Respiratory epithelium is one of the major cell components in human lungs and alveolar epithelial cells are most affected by human aging ^3^. AT2 cells, surfactant secreting cells, serve as the epithelial lineage progenitors in the alveoli ^5^. In order to examine the aging process of human alveolar epithelial cells, AT2 cells were extracted and clustered with purity confirmed (Figure S3A). Differential expression analysis revealed a unique gene expression profile of AT2 cells from each age group (Figure 2A). Notably, aged AT2 cells underwent a significant decrease in cell proliferation rate, as indicated by a reduction in the proportion of cells that were positive for MKI67, HMMR, or TOP2A (Figure 2B). Furthermore, elevated cellular senescence, a hallmark of aging ^36^, was found in the aged AT2 cells. This was confirmed by both bioinformatically increased senescence marker genes (Figures S3B-C) and cellular senescence score returned by the average expression of core senescence genes as described previously ^31^ (Figure 2C), and histologically upregulated P21 (CDKN1A) protein expression, a commonly recognized cellular senescence marker, in HTII-280+ AT2 cells in human lung sections from aged donor (Figure 2D).

**Figure 2.**
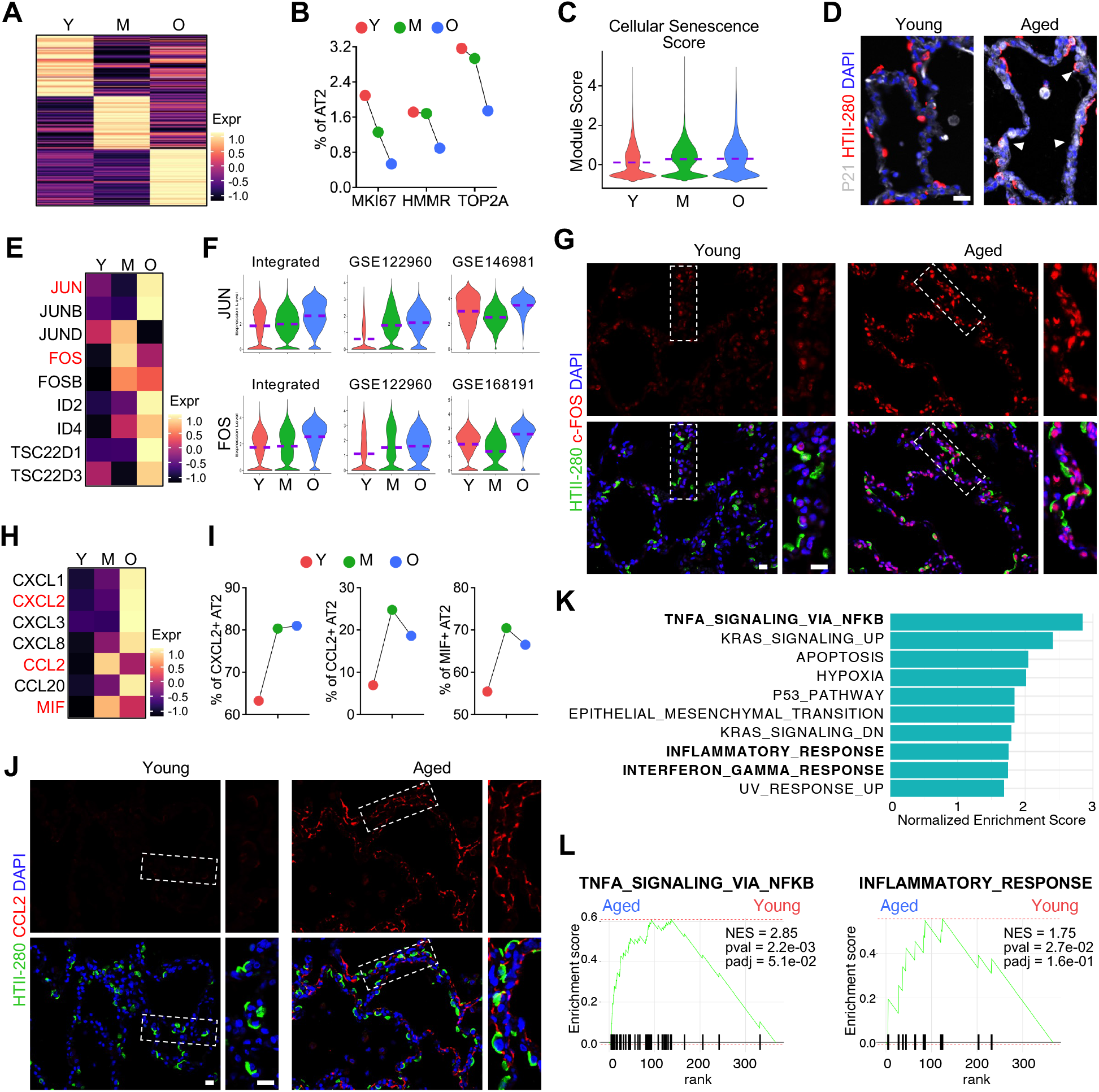
Elevated cellular senescence and inflammation in AT2 cells of aged human lungs. **(A)** Heatmaps showed the expression of top 100 genes (rows) with distinct gene profiles of AT2 cells from subjects of different age stages (columns). **(B)** Percentages of AT2 cells positive for cell proliferating marker genes, MKI67, HMMR, and TOP2A, were quantified from subjects of different age stages. **(C)** Violin plots showed the cellular senescence scores of AT2 cells from different age stages. The cellular senescence score was defined as the average transcription of 130 core senescence genes from CSGene that were detectable in the AT2 cells. **(D)** Representative immunofluorescence staining of cell senescence marker (CDKN1A/P21) and human AT2 cell marker (HTII-280) on human lung sections from young and aged donors. White arrowheads indicated the senescent AT2 cells. **(E** and **F)** Heatmap (**E**) and violin plot **(F**) visualization of the upregulated transcriptional factor genes in aged AT2 cells in the integrated data and representative individual datasets. **(G)** Immunofluorescence staining presented the protein levels of representative AP-1 transcription factor gene, c-FOS/FOS, in AT2 cells (HTII-280+) on human lung sections from young and aged donors. Boxed regions were magnified. **(H)** Heatmaps showed the transcriptional levels of chemokine/cytokine genes in aged AT2 cells in the integrated data. **(I)** Percentage quantifications of AT2 cells positive for the representative chemokine/cytokine genes, CXCL2/CCL2/MIF, in lungs from subjects of different age stages. **(J)** Immunofluorescence staining for the representative chemokine protein, CCL2, in alveolar regions on human lung sections from young and aged donors. Boxed regions were magnified. **(K** and **L)** Top canonical Hallmark pathways from GSEA (pval < 0.05) on genes upregulated in aged AT2 cells identified by Fast Gene Set Enrichment Analysis (FGSEA) were illustrated (**K**) and representative pathways were visualized by enrichment plots (**L**). Purple dashed lines in the violin plots indicated the mean levels of module score and gene transcriptions. Y, young donors (Age<40), M, middle-aged donors (Age40-60), O, aged (old) donors (Age>60). Scale bar: 20 µm (**D**, **G**, and **J**).

Expression of surfactant protein genes and lamellar body marker gene ABCA3 represents the normal functions of AT2 cells ^37^ and these genes showed comparable transcription in the AT2 cells of different age stages, while ETV5, a transcription factor essential for the maintenance of AT2 cells ^38^, showed slightly defected transcription in aged AT2 cells in most individual datasets (Figure S3D). Cigarette smoking can induce some deleterious effects on alveolar epithelial cells ^39^. To study the effects of smoking on human AT2 cells, we introduced the identity of smoking history into the differential expression analysis of human AT2 cells (Figure S3E). The surfactant protein genes and ETV5 were consistently downregulated in smoker AT2 cells in the integrated data and most individual datasets (Figure S3E).

The other downregulated genes in aged AT2s were represented by Small Nucleolar RNA Host Genes (SNHG7 and SNHG8), Keratin genes (KRT7, KRT8, KRT18, and KRT19), S100 Protein genes (S100A10, S100A14, and S100A16), Serpin family genes (SERPINA1, SERPINB1, and SERPINH1), and Solute carrier family genes (SLC1A5, SLC25A3, SLC25A5, SLC25A6, SLC34A2, and SLC39A8) (Figures S4A-B). Loss of functions of these representative genes in aged AT2 cells were further confirmed by violin plots in the integrated data and representative individual datasets (Figure S4C). Among these downregulated genes, those representing the epithelial signatures were dominant (Figure S4B and C). Defected transcriptions of these genes, as well as other downregulated epithelial identity genes (Figure S4D) and declined epithelial identity score returned by the average expression of these genes (Figure S4E), suggested a loss of the epithelial identities in the aged AT2 cells in human lungs. To confirm the decreased gene transcriptions in aged AT2 cells, KRT8, KRT19, S100A14, and SLC25A3 were chosen as the representative verifications on the young and aged human lung sections (Figures S4F and G). Although the specific and dysregulated transcriptions of these genes in AT2 were identified here, their functions are largely unknown. Among them, we reported recently that SLC39A8 (ZIP8) is required for AT2 cell renewal in murine and human lungs ^14^.

### Aberrantly elevated AP-1 transcription and cellular inflammation in human AT2s of aged lungs

Surprisingly, a set of Activator Protein-1 (AP-1) transcription factor subunit genes, including JUN, JUNB, JUND, FOS, and FOSB, were found elevated both bioinformatically (Figures 2E-F) and histologically (Figures 2G and S5A) in the aged AT2 cells. AP-1 has been demonstrated to lead a transcription factor network that drives the transcriptional program of senescent cells ^40^, implying a relevance of these gene activations with AT2 cell senescence in our study. The other significantly activated transcription factors genes included ID genes (ID2 and ID4) and TSC22 domain family genes (TSC22D1 and TSC22D3) (Figures 2E), the roles of which in AT2 cells are to be determined. Alveolar epithelial cells have been reported to secrete chemokines in response to some pathogenetic stimulations ^41,42^. In the aged AT2 cells, several chemokine and cytokine genes, such as CXCL1, CXCL2, CXCL3, CXCL8, CCL2, CCL20, and MIF (macrophage migration inhibitory factor), were found to be remarkably activated (Figures 2H and S5B). Among these genes, CXCL2, CCL2, and MIF positive AT2 cells were representatively quantified and were found to be increased with aging (Figure 2I), although their receptors were lowly transcribed in the integrated data (Figure S5C). CCL2 was chosen as the representative gene to be histologically verified in the age AT2 cells on human sections (Figure 2J). To eliminate the effect of smoking, one of the common external stimuli to induce ectopic epithelial inflammatory ^43^, the transcriptions of these chemokine genes were viewed separately regarding to the smoking histories of the subjects. These genes were found to be consistently upregulated in aged AT2 cells from both non-smoker and smoker subjects (Figure S5B). These secreted chemokines and cytokines suggested activated cell-cell communications of aged AT2 with the alveolar microenvironments, although most of their receptor genes were of low transcriptional levels except CD74 (Figure S5C), the canonical receptor for MIF ^44^.

To study the potential pathways regulating the DE genes of the aged AT2 cells, we employed an R based fast gene-set enrichment analysis (FGSEA) for the pathway analysis on the aged vs young AT2 cells. Remarkably, among the top hallmark pathways of aged AT2 cells many are related to activated inflammatory response (Figures 2K and L). To further confirm this, we input the significantly upregulated genes of aged AT2 cells from the integrated data and representative individual datasets into another leading pathway analysis application, Ingenuity Pathway Analysis (IPA). Consistently, most of the top activated signaling pathways were myeloid and lymphoid cell related inflammation pathways (Figure S6A-E). These data suggested an activated inflammatory reaction in aged human AT2 cells, potentially due to a long-term exposure of these cells to the stimuli, such as air pollutants and microbes, from the external environments.

Another major alveolar epithelial component, AT1s, was also extracted from the integrated data (Figure S7A). Similar to aged AT2s, the aged AT1 cells exhibited an increased cellular senescence (Figures S7B-D). Besides the epithelial identity genes (EPCAM, NKX2-1, KRT8, and KRT19), other genes associated with AT1 spreading (AGER), development and differentiation (HOPX and LY6E), and normal functions (PDPN, CLDN18, and CAV1) were also found to be defectively expressed in aged AT1s (Figures S7E and F), indicating a set of defects in aged AT1s.

The human lung airway epithelium is composed of several known epithelial lineages, in which the ciliated cells were most abundant and functionally distinct. Here we extracted the ciliated cell clusters and identified the differentially expressed genes at different age stages (Figure S8). Of these genes, the critical transcription factor genes, Cilia/Flagella/Tubulin/Dynein/Mucin genes, Coiled-coil domain containing family genes, and some open reading frame (orf) transcripts were specific to ciliated cells (Figure S8B) and were dysregulated in aged ciliated cells (Figure S8A), although the roles of some of genes are mostly unknown.

### Diminished Collagen and Elastin gene transcription in aged lung mesenchyme

Aging has been considered as the driving force of the progressive interstitial lung diseases such as idiopathic pulmonary fibrosis (IPF), a disease with accumulated fibroblasts and excessive extracellular matrix (ECM) production as the hallmarks ^45^. To comparatively profile the fibroblasts from lungs of donors at different ages, the mesenchymal cells were extracted from the integrated data and re-clustered (Figure S9A). Intriguingly, a declined ECM transcriptional program was observed in the aged fibroblasts (Figure 3A). More specifically, decreased transcriptions of matrix genes including COL1A1, COL1A2, COL3A1, COL6A1, COL6A2, FN1, and ACTA2 (Figure 3B), as well as a reduced level of Matrix gene score, defined by the average expression of these genes, were identified in the aged fibroblasts (Figure 3B). This observation was even more obvious in the individual databases, especially in the datasets with considerable numbers of fibroblasts (Figure S9B). To eliminate the potential bias caused by the smoking history of the subjects, the fibroblasts from lungs of non-smokers and smokers were examined separately and downregulated collagen gene expression was observed in aged fibroblasts from both non-smokers (Figure S9C) and smokers (Figure S9D). These observations were verified by the immunostainings for Collagen I, α-SMA (Figures 3D) and Collagen VI (Figure S9E) on lung sections from young and aged donors, and their protein intensity quantifications (Figure 3F).

**Figure 3.**
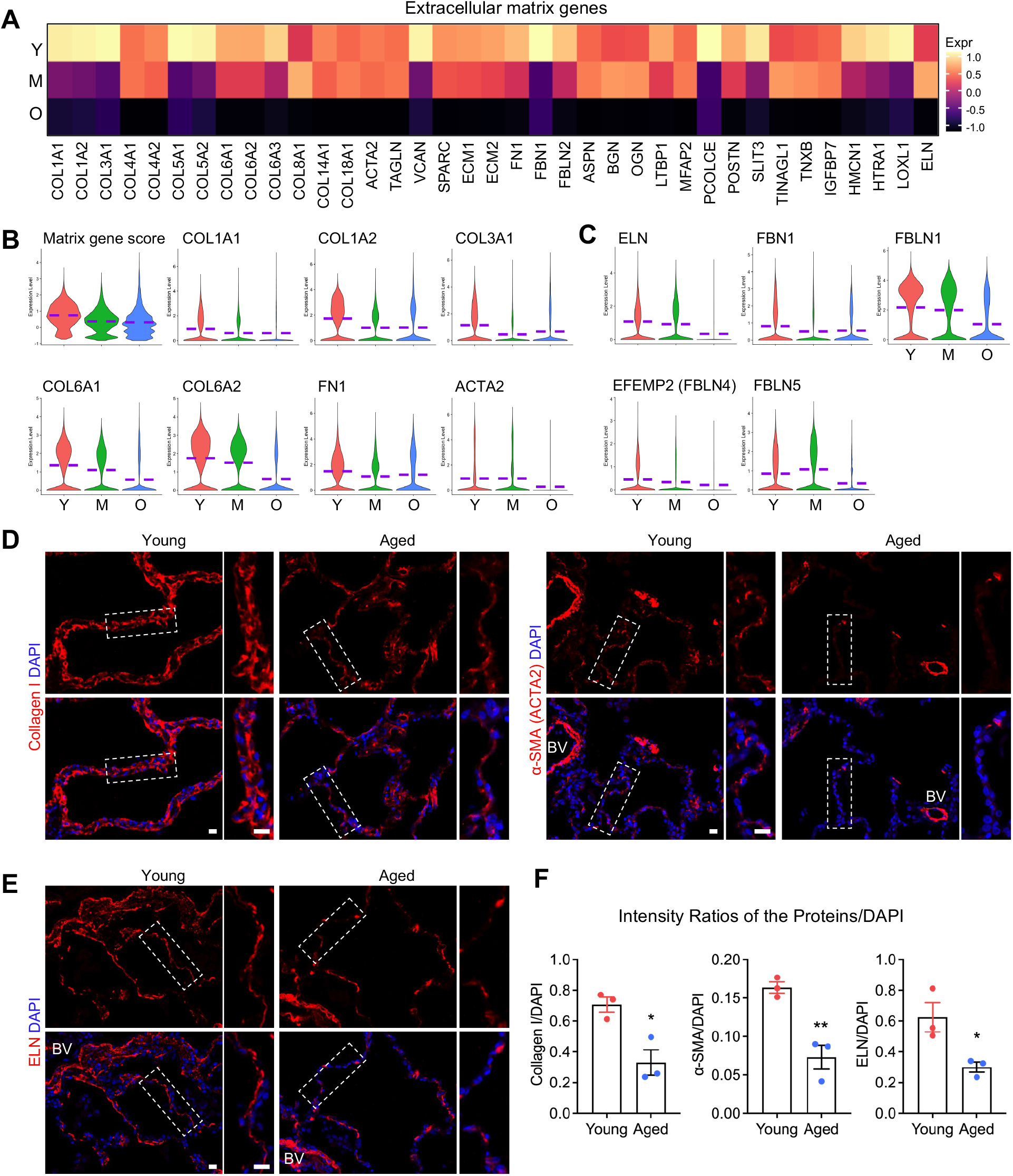
Decreased extracellular matrix gene transcription in aged lung mesenchymal cells. **(A)** Heatmaps showed the expression of the extracellular matrix genes in the mesenchymal cells from subjects of different age stages. **(B)** The Matrix gene score and transcriptions of representative collagen genes in mesenchymal cells from subjects of different age stages were visualized by violin plots. The Matrix gene score was defined as the average transcription of COL1A1, COL1A2, COL3A1, COL6A1, COL6A2, FN1, and ACTA2. **(C)** Violin plot showed the transcriptions of genes specific to elastic fiber components and assembly in mesenchymal cells from subjects of different age stages in the integrated data. **(D-E)** Immunofluorescence staining for Collagen I, α-SMA (ACTA2), and ELN (Elastin) on human lung sections from young and aged donors. Boxed regions were magnified. BV, blood vessel. **(F)** Quantified intensity ratios for Collagen I, α-SMA (ACTA2), and ELN (Elastin) staining compared to nuclear DAPI on human lung sections from young and aged donors. Lung sections from three young and three aged donors were examined (n = 3 per group). Purple dashed lines in the violin plots indicated the mean levels of gene transcriptions. Y, young donors (Age<40), M, middle-aged donors (Age40-60), O, aged (old) donors (Age>60). Scale bar: 20 µm (**D** and **E**). * p < 0.05, ** p < 0.005.

The synthesis and function of Elastin (ELN), as a unique ECM component with a specific function to provide elasticity (both compliance and recoil) to the lungs, are rapidly declined with age ^46^. As one of the decreased ECM genes in aged fibroblasts (Figure 3A), the transcription of Elastin, as well as other elastic fiber component and assembly genes, were found even more consistently downregulated in both the integrated data (Figure 3C) and most individual datasets (Figure S10). Consistent with the RNA-levels in the scRNA-seq datasets, the protein level of ELN (Elastin) in aged lung sections was significantly declined (Figures 3E and F). This observation suggested an obvious decrease in the elasticity of aged human lungs.

### Dysregulated transcriptional profile and aberrantly activated inflammation in aged lung macrophages

Pulmonary immune homeostasis in human lungs is maintained by a complex network of immunocytes derived from myeloid and lymphoid lineages ^47^. To profile the immunocytes from human lungs of different age stages, we extracted the myeloid and lymphoid lineages identified above (Figure 1D). Macrophages, the largest myeloid population in the integrated data, was extracted and re-clustered (Supplementary Figure S11A). Comparative analysis revealed dramatically differentially expressed gene profiles between the young and aged lung macrophages (Figure S11B). Many of the DE genes were functional genes specific to macrophages (Figure S11C). The downregulated genes included several gene families, such as LAIRs (Leukocyte associated immunoglobulin like receptor genes), CTSs (Cathepsin genes), CSTs (Cystatin genes), SH3BGRL (SH3 domain binding glutamate rich protein like genes), MS4As (Membrane spanning 4-domains genes), and CD68 (Figures S11B and C). All these genes have shown some relevance with either inflammation inhibition or M2 macrophage polarization, suggesting an aberrant age-associated decay in anti-inflammatory of the lung macrophages from aged subjects. Together with these data, ectopic activation of macrophage-mediated inflammatory genes (MSR1, ALOX5, and SCD) and macrophage-specific antigen presentation genes (HLA-DQA1, HLA-DRB5, and HLA-DRB6) (Figures S11B and C) further revealed an activated inflammatory genetic program modulated by aged macrophages in human lungs. Remarkably, the elevated genes in aged lung macrophages included a set of macrophage specific chemokine genes, such as CXCL2, CXCL3, CXCL5, CXCL8, CXCL16, CCL12, CCL20 (Figure 4A), and the transcriptions of these chemokine genes were consistently increased in the age macrophages of most individual datasets (Figure 4B). Many of these genes, as well as other chemokine and cytokine genes, were found to be extremely highly expressed in the lung macrophages from smoking subjects (Figures 4C and D), indicating a positive correlation between smoking and macrophage-associated lung inflammation.

**Figure 4.**
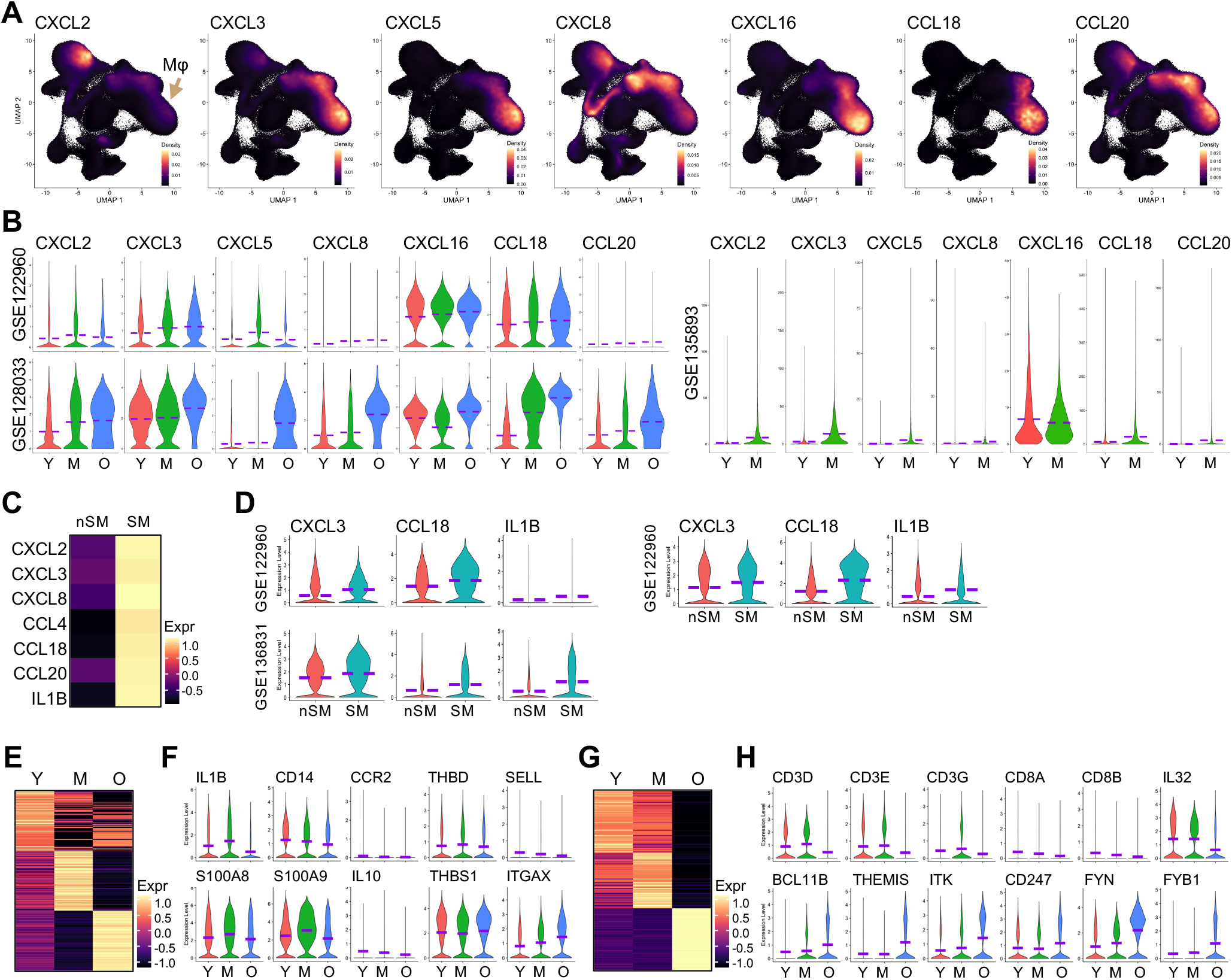
Hyperactivated chemokine expression in macrophages from aged and smoking lungs. **(A** and **B)** Density plots from the integrated dataset (**A**) showed the specification and violin plots from the representative individual datasets (**B**) showed the upregulation of macrophages specific chemokine genes in lungs from aged subjects. **(C)** Heatmaps showed the relative transcription levels of chemokine/cytokine and receptor genes in lung macrophages from smoking and non-smoking subjects. **(D)** Elevated transcriptions of representative chemokine/cytokine genes, CXCL3, CCL18, and IL1B, in macrophages from smoking subjects were visualized by violin plots in individual datasets. **(E** and **G)** Differential analysis revealed the top 100 differentially expressed genes (rows, ranked by log fold-change of the average expression) in lung monocytes (**E**) and T cells (**G**) from lungs of different age stages. **(F** and **H)** Violin plots showed top dysregulation of genes in lung monocytes (**F**) and T cells (**H**) from aged subjects. Most of the genes were downregulated in aged monocytes. Purple lines in the violin plots indicated the mean levels of gene transcriptions. Purple dashed lines in the violin plots indicated the mean levels of gene transcriptions. Y, young donors (Age<40), M, middle-aged donors (Age40-60), O, aged (old) donors (Age>60). nSM, non-smoker, SM, smoker.

At least two distinct macrophage populations exist in human lungs at homoeostasis characterized by their unique locations, properties, and functions: alveolar and interstitial macrophages ^48^. By checking the transcriptions of their canonical signature genes (Figure S11D) ^49^, the alveolar and interstitial macrophage populations were respectively identified in the macrophage dataset (Figure S11E). Comparative analysis returned the transcriptions of chemokine, cytokine, transcriptional factor, and enzymatic genes specific to alveolar macrophage and the relatively high expression of MHC class II antigen-presenting protein, FABP protein, and Complement C1q protein genes in interstitial macrophages (Figure S11F). We also demonstrated the transcriptional factor genes differentially expressed in the alveolar and interstitial macrophages (Figure S11G), and the functional involvements of these genes in lung macrophages are largely undetermined. CD74, the canonical receptor for epithelial cell specific cytokine MIF (Figure S5C), was intriguingly found to be predominantly expressed in interstitial macrophages (Figures S11H and I), and its transcriptional level in lung macrophages was significantly augmented with age (Figures S11H and J), implying an increasing cell interaction between epithelial cells and macrophages with the aging of lungs.

Another lung myeloid lineage, monocyte, is capable to differentiate into lung macrophage ^49^ and has been identified with specific inflammatory gene expression signatures in the integrated data (Figures 1D and S12A). Lung monocytes have been suggested to show a reciprocal and interdependent relationship with lung neutrophils during early lung inflammatory response ^50^. Although the transcriptions of neutrophil specific genes such as S100A8 and S100A9 were detected in the monocyte cluster (Figure S12A), we could not identify a separated neutrophil population. Comparative analysis suggested a dysregulated gene profiles in aged monocytes with most of the monocyte specific inflammatory genes downregulated (Figures 4E and F). Similarly, a distinct DC cluster was identified in the integrated data and most of the DC specific antigen-presenting and cell surface receptor genes were downregulated in the aged DC cells (Figure S12B and C).

We have identified the lymphoid lineages in the integrated data (Figure 1D) and these lineage clusters were extracted together with a jointed small cluster of master cells (Figures S13A and B). Differential expression analysis of different lineages revealed distinct transcriptional patterns of each cell type (Figure S13C). Comparative analysis suggested that the gene expression profiles of cells from young and middle-aged donors were more comparable and were jointly more distinct to those from aged donors (Figures 4G, S13D, F, H, and J). Specifically, the genes of TCR (T cell receptor) complex (CD3D, CD3E, and CD3G) and co-receptors (CD8A and CD8B), as well as the T cell specific TNFα inducing proteins (IL32), were consistently decreased in the T cells from aged subjects (Figures 4G and H). On the other hand, the upregulated genes in aged T cells included genes associated with T cell differentiation, such as BCL11B and THEMIS, and genes of T cell specific tyrosine kinases, such as ITK, FYN, and FYB1 (Figures 4G and H). Natural killer (NK) cells, as effector lymphocytes of the innate immune controlling several types of microbial infections, were also dysregulated in the aged human lungs (Figures S13D and E), visualized by the defected gene transcriptions of antimicrobial cytotoxicity proteins (GNLY, NKG7, FCGR3A, and CST7), Killer cell lectin like receptor proteins (KLRB1, KLRD1, and KLRF1), granzymes (GZMA and GZMB), chemokines (CCL4, CCL3, and CCL5), and other cell growth and signal-transducing molecules (CD7 and FGFBP2), suggesting a decay in the effective regulation of cell cytotoxicity in the aged NK cells.

The other innate lymphoid cell lineages identified in the human lungs were the B cells and their progenies, plasma cells, both of which were responsible primarily for the basic functions of antibody production ^51^. Dysregulated gene expression profiles were observed in both aged B and plasma cells (Figures S13F-I). In aged B cells, genes associated with B cell development, survival, and maturation (CD79A/B, TNFRSF13C, CCR7, and LY9) as well as with immunoglobulin production (IGHM), were downregulated, while genes regulating B cell differentiation (AFF3, EBF1, PAX5, BCL11A, and BLK) and signaling (BANK1 and MS4A1) were selectively elevated (Figures S13G). Plasma cells represent the terminal differentiation step of mature B lymphocytes characterized by large amount of antibody secretion ^52^, however in the plasma cells from age subjects, the transcription levels of immunoglobulin genes were consistently declined (Figures S13H and I), suggesting a decay in immune response in aged human lungs.

New insights and perspectives have been disclosed recent years on mast cells ^53^, another myeloid lineage but with closer signatures with lymphoid lineages (Figures 1D and S13A-B). Genes regulating mast cell activation, survival, and differentiation were mostly upregulated (IL1RL1, MS4A2, KIT, and CPA3) although a few downregulated (GATA2, TPSB2, and TPSAB1) in the aged mast cells (Figures S13J and K), implying an abnormal mast cell regulating program in aged human lungs.

### Hypofunctions of endothelial cells in aged human lungs

Pulmonary endothelial cells are an essential component for the gas exchange and pulmonary circulation machinery of the human lungs. To profile the lung endothelium, the endothelial cell clusters were extracted from the integrated data (Figure S14A). It is known that cigarette smoking has systemic effects in altering lung functions, one of the major effects is vascular damage and endothelial dysfunction ^54^. As expected, the lung endothelial cells from smoking subjects exhibited defected transcriptions of genes that were critical for endothelial cell signal transduction (KDR), survival (TIE1), development (EFNA1, EFNB2, RHOA, and TRIOBP), and junctional integrity (PECAM1 and CDH5) (Figure 5A). Lung endothelial cells have been proposed to be heterogenous ^55,56^, and vascular and lymphatic endothelial cell clusters were distinctly identified based on their signatures (Figure 5B and C). Notably, most of the canonical genes related to endothelial angiogenesis (VWF, TIE1, and CAV1), integrity (CDH5 and CLDN5), and differentiation (TMEM100, SOX18, SOX17, AQP1, KLF2, and EGFL7) showed blunted transcriptions in aged vascular endothelial cells, while most of the top elevated genes in aged vascular endothelial cells were protein kinase and phosphatase genes (Figures 5D, E, and S14B). Similarly, the functional genes associated with lymphatic vessel growth (FLT4, TM4SF18, and MMRN1), development (TBX1, PROX1, and PDPN) and cell transmigration (CCL21 and LYVE1) were dramatically downregulated and multiple kinase and phosphatase genes were aberrantly activated in the aged lymphatic endothelial cells (Figures S15A-C). Although the major metabolic processes including oxidative phosphorylation have been reported in stimulated endothelial cells ^57^, the mechanisms underlying were largely unclear. To study the pathways regulating kinase and phosphatase gene activation in aged endothelial cells, we run the IPA analysis on the upregulated genes in the aged vascular and lymphatic endothelial cells respectively. The top signaling pathways of both analyses included activated protein kinase A signaling and integrin signaling, as well as inhibited adherens junction signaling and PTEN signaling (Figures S15D and E), suggesting a dysregulated protein phosphorylation and dephosphorylation program in the vascular and lymphatic endothelial cells in aged human lungs.

**Figure 5.**
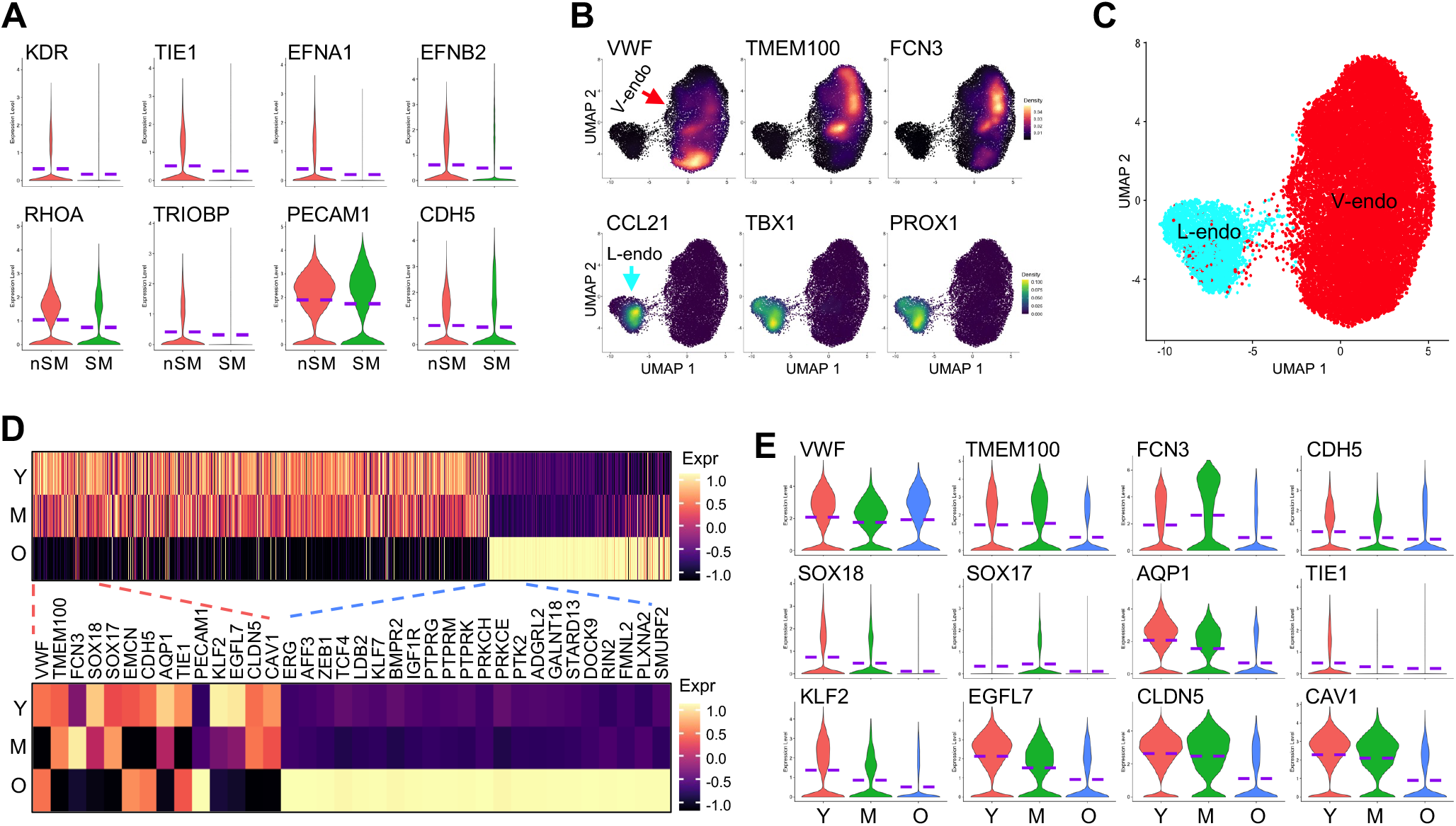
Hypofunctions of vascular endothelial cells in lungs from aged donors. **(A)** Violin plots presented the transcriptions of genes related to vascular formation and barrier integrity in the total endothelial cells from non-smokers and smokers. nSM, non-smoker, SM, smoker. **(B)** Density plot visualization of the relative expression of vascular and lymphatic endothelial specific marker genes. Red and cyan arrows indicated the V- and L-endothelial cell clusters, respectively. **(C)** UMAP showed the identification of lung vascular and lymphatic endothelial cells isolated from the integrated data. **(D)** Heatmap visualization of the top differentially expressed genes (columns, ranked by log fold-change of the average expression) in the vascular endothelial cells from donors at different age stages (rows). **(E)** Transcriptions of the representative downregulated functional genes in vascular endothelial cells from lungs of aged subjects. Purple dashed lines in the violin plots indicated the mean levels of gene transcriptions. V-endo, vascular endothelial cells, L-endo, lymphatic endothelial cells. Y, young donors (Age<40), M, middle-aged donors (Age40-60), O, aged (old) donors (Age>60).

### Constantly activated cell-cell communications between AT2 and macrophages in aged lungs

Qualitative deficits during aging are believed to lead to abnormal cell-cell communication mechanisms in lungs ^3^. To determine the alterations in cell-cell communications during aging, we employed the CellChat, a R toolkit for inference, visualization, and analysis of cell-cell communication ^58^, and elucidated a dynamic change in cell-cell communications with the aging of lungs (Figure S16A). We noticed an increased cell-cell communication profile in AT2 cells and macrophages in aged lungs when the interaction weights among other cell types were mostly compromised. The increased cell-cell communication was even more obvious when AT2s were set as the sender cells (Figure 6A). Visualizations of all the individual L-R (Ligand-Receptor) pairs emanated from AT2s suggested that the dominant significant interaction in the aged lungs was MIF and its receptor CD74 (and CD44, CXCR4) (Figure S16B). The interaction strength of MIF-CD74 among cell types was more even in lungs of young subjects but became more condensed between AT2s and macrophages in the lungs from aged subjects (Figure 6B). Heatmap quantification further demonstrated that in the aged lungs the AT2s were the only dominant originator cells of the MIF signaling pathway network (Figure 6C). This may be explained by the more dominant and elevated expression of MIF in AT2s and CD74 in macrophages in aged lungs (Figures 2H, I, S11J, and S16C). Together, with aberrant MIF-CD74 signaling pathway in aged AT2s and macrophages as an example, we established a dysregulated cell-cell interactome in aged human lungs that may interlink the exhausted cellular niches with compromised supportiveness among alveolar cell types and subsequent accumulated cellular deficits along with aging.

**Figure 6.**
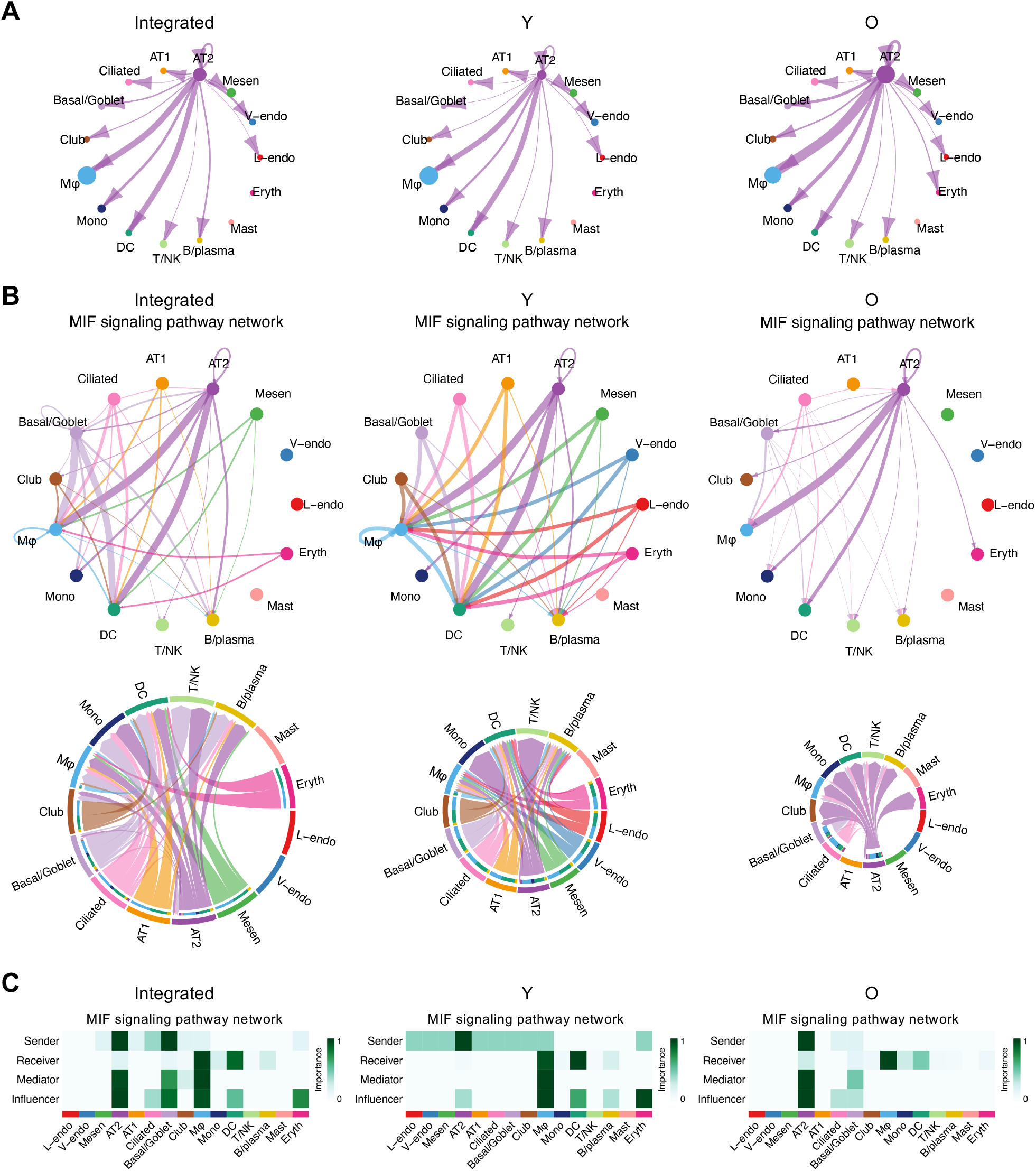
Constant cell-cell communications between AT2 and macrophages in aged lungs. **(A)** Overall views of inferred communication edge weights of all ligand-receptor interactions sent from AT2 cells in the integrated, young (Y) and aged (O) datasets. The communications between AT2-macrophage became stronger in aged lungs. **(B)** Circle plot (top panels) and Chord diagram (bottom panels) visualization of cell-cell communications by MIF signaling pathway network among cell types in the integrated, young (Y) and aged (O) datasets. The communications between AT2-macrophage were strong and were stronger in aged lungs. **(C)** Identification of dominant senders, receivers, mediators and influencers in the intercellular communication network by computing MIF signaling pathway network centrality measures for each cell type in the integrated, young (Y) and aged (O) datasets. The communications by MIF signaling pathway network were wide with several cell types as senders in young lungs but were more specific to AT2-macrophage with AT2 as the only sender in the age lungs. L/V-endo, lymphatic/vascular endothelial cells, Mesen, mesenchymal cells, AT1/2, alveolar type I/II epithelial cells, Macro, macrophages, Mono, monocytes, DC, dendritic cells, Eryth, erythrocytes. Y, young donors (Age<40), M, middle-aged donors (Age40-60), O, aged (old) donors (Age>60).

**Figure 7.**
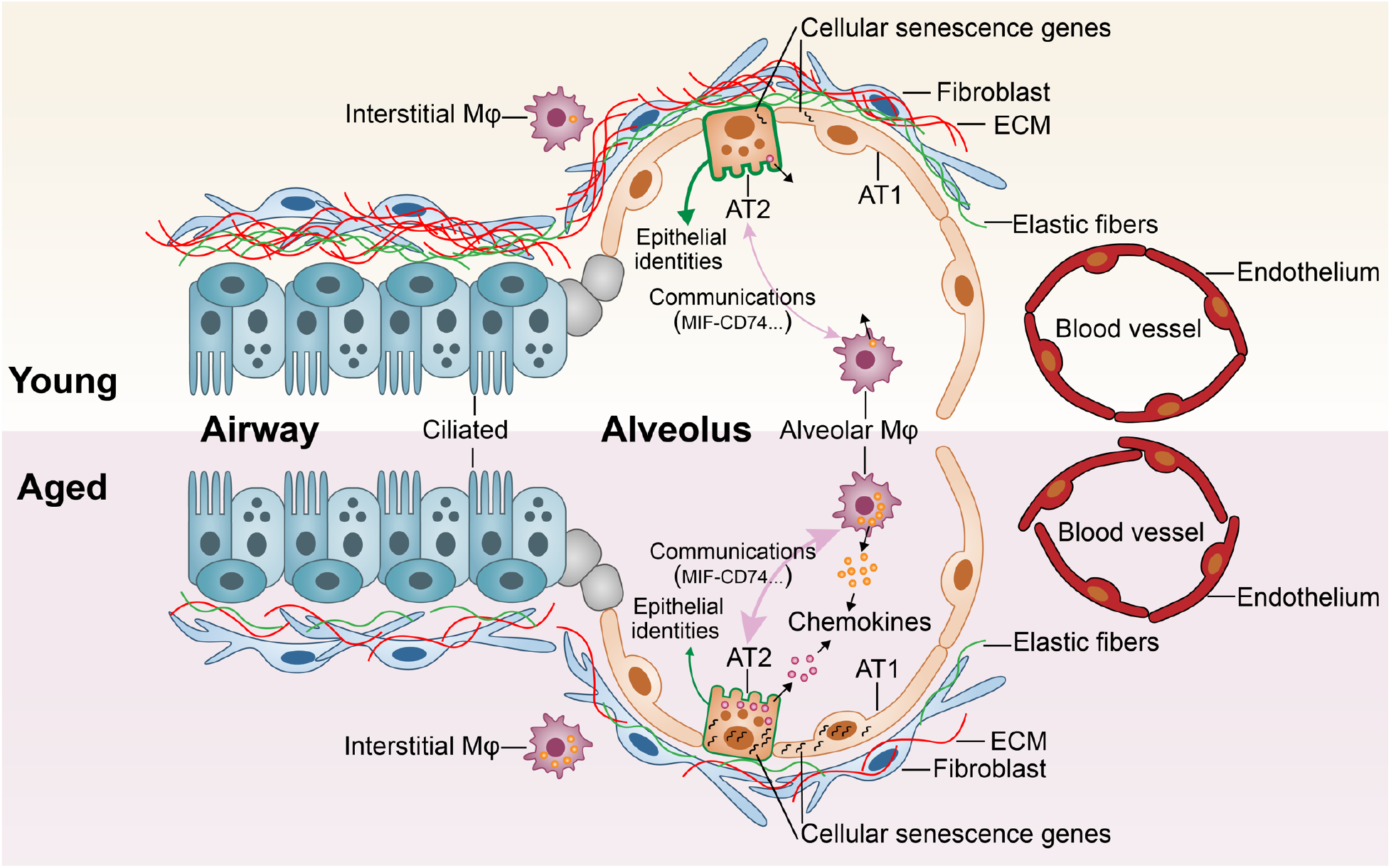
Schematic summarization of the histopathological, cellular, and molecular alterations in aged human lung. The aged lungs have a general aged-associated decay in multiple cell types, suggesting a general compromised lung function. AT1/2, alveolar type I/II epithelial cells, Mφ, macrophages, ECM, extracellular matrix. The basement membranes are represented by elastic fibers.

Altogether, we have provided a comprehensive single cell multi-omics atlas of healthy human lungs with distinct cellular lineages covering all age ranges. The gradual dysregulation of the genetic profiles in the major cellular lineages proposes an increasing general hypofunction of human lungs with age, highlighting a significant decline of the AT2 niche. Dysregulated alveolar niches were further deteriorated by impaired macrophages and endothelial cells in aged lungs. The proposition raises a hypothesis that the higher risk of human lung diseases in aged populations may result from the accumulations of multiple cell lineage exhaustions and the miscommunications among these defected cell lineages. This study provides a comprehensive portrait of the healthy human lungs. The identification and detailed description of the cell-type/lineage specific transcriptional profiles provides a highly accessible reference to explore the aged-related cell deficits in human lung diseases.

## DISCUSSION

As the rapid increase of aging population worldwide, degenerative changes in lungs have been found to be strongly correlated with the development and incidence of chronic respiratory diseases, such as COPD, IPF, and lung cancers. The cellular and molecular hallmarks of aging, including genomic instability, telomere attrition, epigenetic alterations, loss of proteostasis, deregulated nutrient sensing, mitochondrial dysfunction, cellular senescence, stem cell exhaustion, and altered intercellular communication, have been summarized ^36^. A number of transcriptomic studies of murine lung aging have been recently performed ^25,26^, however, a longitudinal transcriptomic profiling of human lungs at single cell level across all adult age ranges is not available. To systemically characterize the effect of aging on the transcriptomic changes of human lungs, we collected and integrated the open-accessed single cell transcriptomic studies on adult human lungs of all age ranges and applied a comprehensive analysis covering all the lung cell lineages. An overall dysregulated genetic program has been demonstrated in the aged lungs with the abnormal transcriptomic changes within all lineages including mesenchymal cells, epithelial cells, endothelial cell, myeloid cells, and lymphoid cells.

The lung epithelium in aging shows deficits in progenitor cells in the airway and alveolar parenchyma. One aging hallmark of the AT2 cells, which are the major progenitor cells in the alveoli, is the depletion of adult stem cell reservoirs and failure in self-renewal and differentiation capacity ^3^. This depletion and dysfunction of AT2 cells may contribute to pulmonary diseases like emphysema and pulmonary fibrosis ^14,31^. However, most of the existing data are derived from rodent studies. Thus, questions remain about how aging reshapes the lung stem niche and how advanced age alters the contributions of epithelial populations to lung regeneration ^3^. In this study we found that the genes driving cellular senescence are elevated, and the genetic programs deriving immune aging such as AP-1 transcription factor subunits are aberrantly activated in AT2 cells in aged lungs. Another consistent feature of both AT2 and AT1 cells from aged lungs is the loss of epithelial identities. These dysregulated genetic programs of aged AT2 cells would trigger some pathogenetic signaling cascades, ultimately leading to defected respiratory epithelium and increased susceptibility to development of chronic diseases. AT2 cells from aged lungs produce higher levels of chemokines, this has been manifested by a previous study on an influenza murine model, in which neutrophils induced by alveolar epithelial cells secreted chemokines increased the mortality of aged animals after infection ^59^. These data propose the effects of aging on pulmonary innate immune dysfunction and the compromised response to invading microorganisms.

The AT2 niche is supported by lung mesenchymal cells, mainly fibroblasts, which have critical roles in lung development, and homeostasis, and lung repair to chronic injuries could contribute to lung diseases. Aging increased the capacities of lung fibroblasts in differentiation into myofibroblasts and restricted their effects in supporting alveolar epithelial renewal ^45^. Increased collagen production in adult fibroblasts is associated with interstitial lung diseases, which have higher morbidity in aged human populations. A most recent comparative analysis of transcriptomic and proteomic data identified increased expression of several ECM gene with age at both RNA and protein levels in specific regions of healthy human lungs ^60^. However, here we reported a consistent defect in collagen gene transcriptions by the total mesenchymal populations from aged human lungs. This inconsistency may rise from either unpurified mesenchymal cell isolation or unbiased selection of lung regions during transcriptomic and proteomic analysis at bulk levels. The mechanisms or consequences of collagen loss in aged lung fibroblasts have rarely been reported, but decreased collagen production in aged skin fibroblasts has been frequently demonstrated to result in fragmentation of the dermal collagen matrix and impairment in the structural integrity of the dermis ^61^. Whether the fibroblasts from these two organs shared similar roles or underlying mechanisms remains to be further investigated. Another observation in the aged lungs was the loss of Elastin (ELN) expression by fibroblasts. Elastin is required for normal lung development and destruction of elastin or abnormality in elastic fiber assembly are the major factors in emphysema and destructive lung diseases. In adults, Elastin conveys elasticity to the lungs and loss of elasticity is associated with reduced lung compliance and recoil leading to an increasing risk of emphysema ^46^. Impaired lung elasticity due to defected Elastin production and elastic fiber assembly might explain the compromised forced vital capacity and expiratory volume in the aged human populations.

The immune response is required to protect against invading pathogens and external material and well-coordinated control of pro-inflammatory and anti-inflammatory mediators is critical to prevent uncontrolled injury and fibrotic deposition ^47^. The lung myeloid cells in conjunction with lymphoid lineages play the central role in orchestrating the pulmonary immune response, however we found a dysregulated genetic program in the immune system of the aged human lungs. Most significantly, the aged lung macrophages exhibited a sustained pro-inflammatory effect and aberrant cytokine gene expression. The dysregulated gene expression profiles in the immunocytes, consistent with recently reviewed ^3^, might reveal an aged-associated decay in the immune function efficiencies in the aged human lungs.

Pulmonary endothelial cells are crucial for gas exchange in the lungs. Endothelial dysfunction increases with age and in chronic respiratory diseases ^62^. However, the roles of endothelial cells in age-related lung impairments are not well studied. Our analysis of the human lung single-cell atlas reveals dysregulated gene expression patterns in both vascular and lymphoid endothelial cell subtypes in aged lungs. These changes reflect deficits in the development, integrity, and maturation of endothelial cells. The involvements of the activated kinase and phosphatase genetic programs in aged endothelia cells are rarely reported. Although some evidence suggested a link between endothelial complications associated with the metabolic syndrome and AMP-activated protein kinase signaling regulated protein phosphorylation ^57^, its connection to aging remains unclear.

Our study is the first to complete the integration of the single-cell transcriptomic data containing all lung cells from healthy subjects covering all age ranges and different sexes and smoking histories. The analyses comprehensively detail the alterations in the age-associated molecular patterns at single cell levels in the human lungs and revealed a systemic decay in lung functions in elderly people. These findings highlight the dysregulation observed in both AT2 stem cells and their supportive niche cells, potentially contributing to the increased susceptibility of aged populations to lung diseases. Although many observations need further functional and mechanism verifications, the current study has drawn a comprehensive transcriptomic map of healthy human lungs, and deep excavation of this map will hopefully improve our understanding of lung degeneration and exhaustion in elder people, as well as age-associated deficiency and pathogenesis in the respiratory diseases.

## STAR METHODS

Detailed methods are provided in the online version of this paper and include the following:

- KEY RESOURCES TABLE
- RESOURCE AVAILABILITY

`o` Lead Contact
`o` Materials Availability
`o` Data and Code Availability
- EXPERIMENTAL MODELS AND SUBJECT DETAILS
- METHODS DETAILS

`o` Data collection
`o` Data integration, visualization, and comparative analysis
`o` Ingenuity Pathway Analysis (IPA)
`o` Fast Gene Set Enrichment Analysis (GSEA)
`o` Inference and analysis of cell-cell communication by CellChat
`o` Histology and immunofluorescence staining
- QUANTIFICATION AND STATISTICAL ANALYSIS

## SUPPLEMENTAL INFORMATION

Supplemental Information can be found online

## ACKNOWLEDGEMENTS

We thank Drs. Tatsuya Tsukui and Dean Sheppard for sharing the metadata of the donors in GSE132771. We thank the lung research community for the open sharing of datasets available. This work was supported by National Institutes of Health grants R35-HL150829, R01-AI052201 (P.W.N.), and P01-HL108793 (P.W.N. and D.J.).

## AUTHOR CONTRIBUTIONS

D.J., and P.N. conceived and supervised the study. X.L., and C.Y. performed the bioinformatic analysis. X.L., X.Z., and J.L. designed and performed the experiments. X.L., X.Z., C.Y., J.L., P.N., and D.J. prepared the figures and wrote the manuscript. All authors reviewed and approved the final version of the manuscript.

## Supplemental information

**Figure S1.**
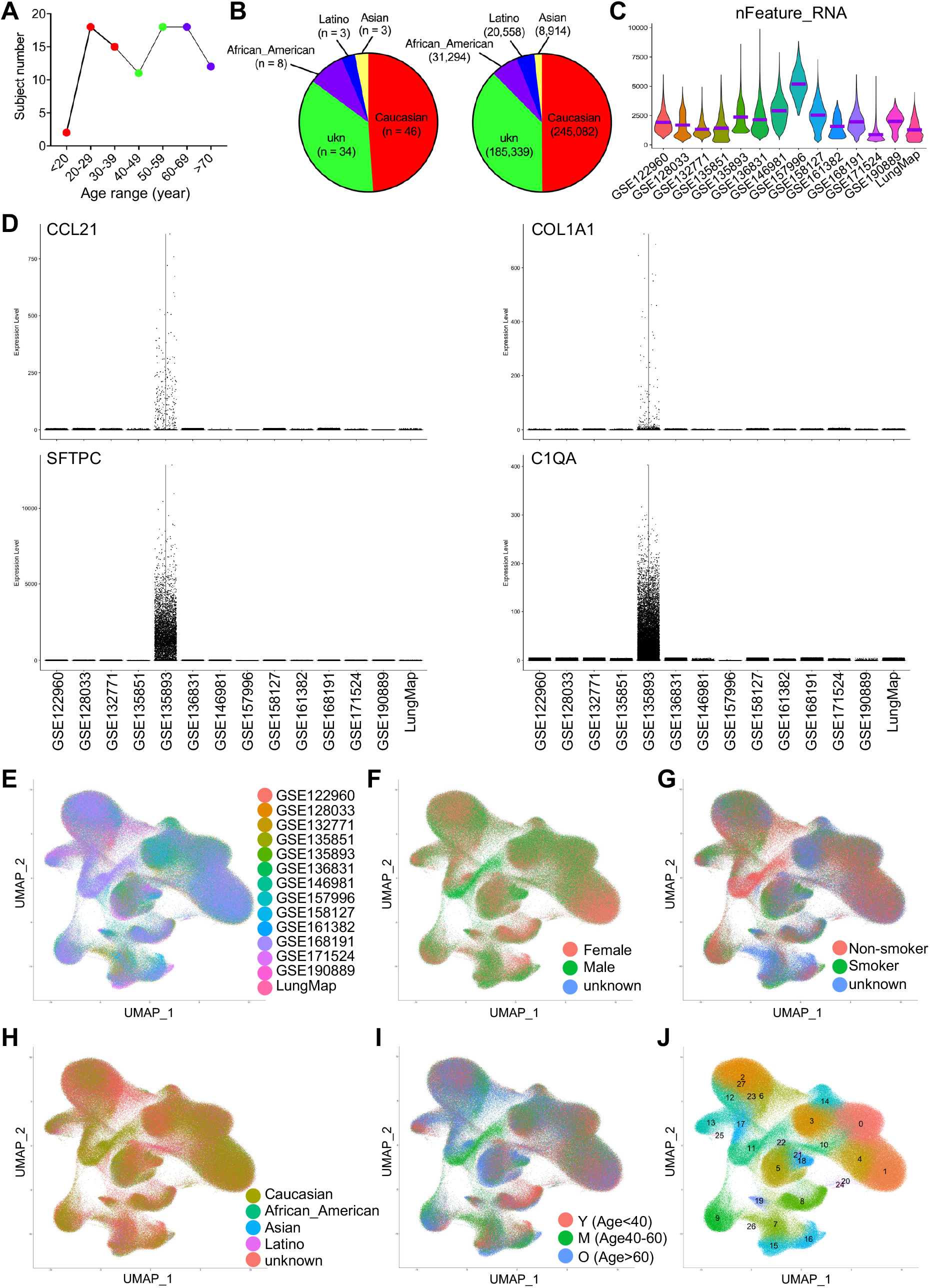
Distributions of cells from subjects of different age stages, datasets, sexes, smoking histories, and races. Related to Figure 1. **(A)** Numbers of subjects from different age ranges indicated a data collection covering a wide age range. **(B)** Subject and cell numbers from donors of different ethnic races were disclosed by pie charts. **(C)** Violin plots showed the mean gene numbers detected (nFeature_RNA) in individual scRNA-seq datasets. Purple lines in the violin plots indicated the mean levels of nFeature_RNA. **(D)** Violin plots of representative cell type specific genes showed different gene-expression scales of GSE135893 dataset with other datasets. **(E-I)** UMAP visualization of cells from different datasets (**E**), sexes (**F**), smoking histories (**G**), ethnic races (**H**), and age stages (**I**) in the integrated data after removing the batch effects. **(J)** 491,187 cells in the integrated data were clustered and 28 major cell clusters were identified. Y, young donors (Age<40), M, middle-aged donors (Age40-60), O, aged donors (Age>60).

**Figure S2.**
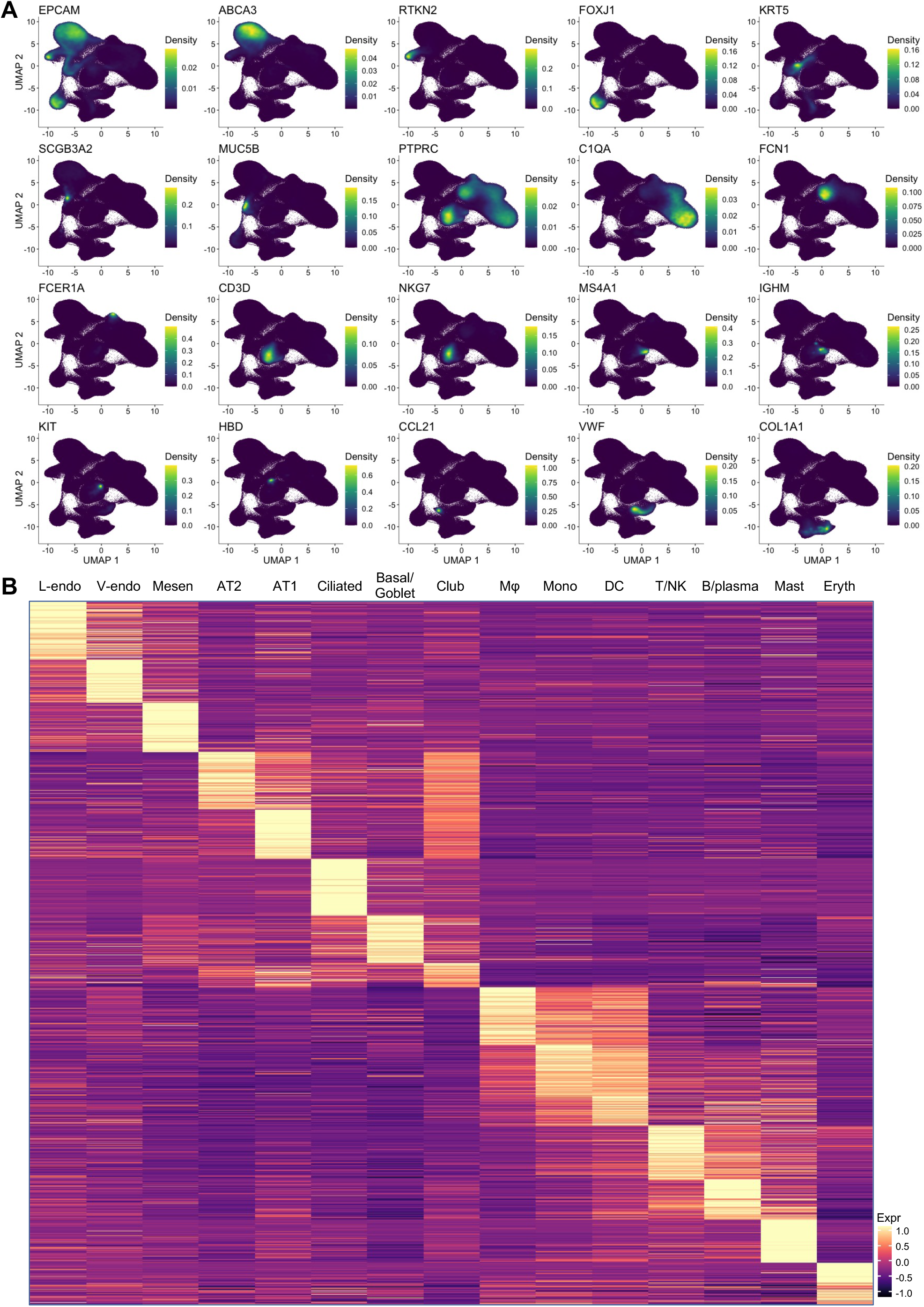
Differential gene expression of each cell type in the integrated data. Related to Figure 1. **(A)** Density plots visualized by UMAPs showed the relative transcriptions of the canonical cell type marker genes in the integrated data. These genes are among the representative genes used for the identification of the major cell type in the integrated data. **(B)** Heatmaps showed the expression of top 100 genes (rows, ranked by log fold-change of the average expression) specific to each cell type (columns) defined in the integrated data.

**Figure S3.**
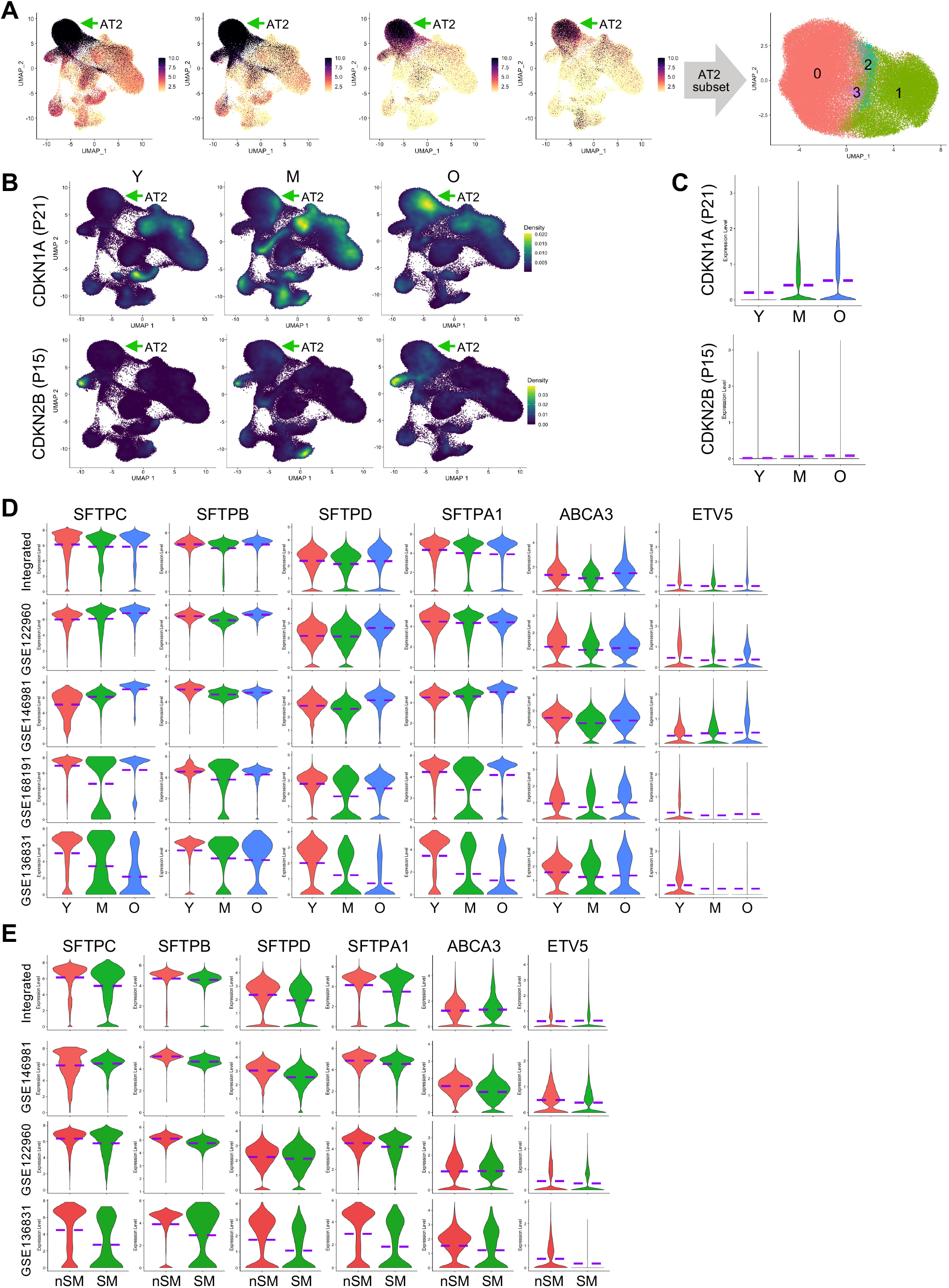
Activated cellular senescence in AT2 cells in aged lungs. Related to Figure 2. **(A)** UMAP visualization of the AT2 subset based on the expression of AT2 specific marker genes. **(B** and **C)** Density plots (**B**) and violin plots (**C**) presented the relative transcriptions of the representative cellular senescence genes, CDKN1A (P21) and CDKN2B (P15), in AT2 from subjects of different age stages. Green arrows indicated the AT2 clusters. **(D** and **E)** Violin plot visualization of surfactant genes and AT2 transcriptional factor gene in AT2 cells from subjects of different age stages (**D**) and smoking histories (**E**) in the integrated and representative individual datasets. Purple dashed lines in the violin plots indicated the mean levels of gene transcriptions. Y, young donors (Age<40), M, middle-aged donors (Age40-60), O, aged (old) donors (Age>60). nSM, non-smoker, SM, smoker.

**Figure S4.**
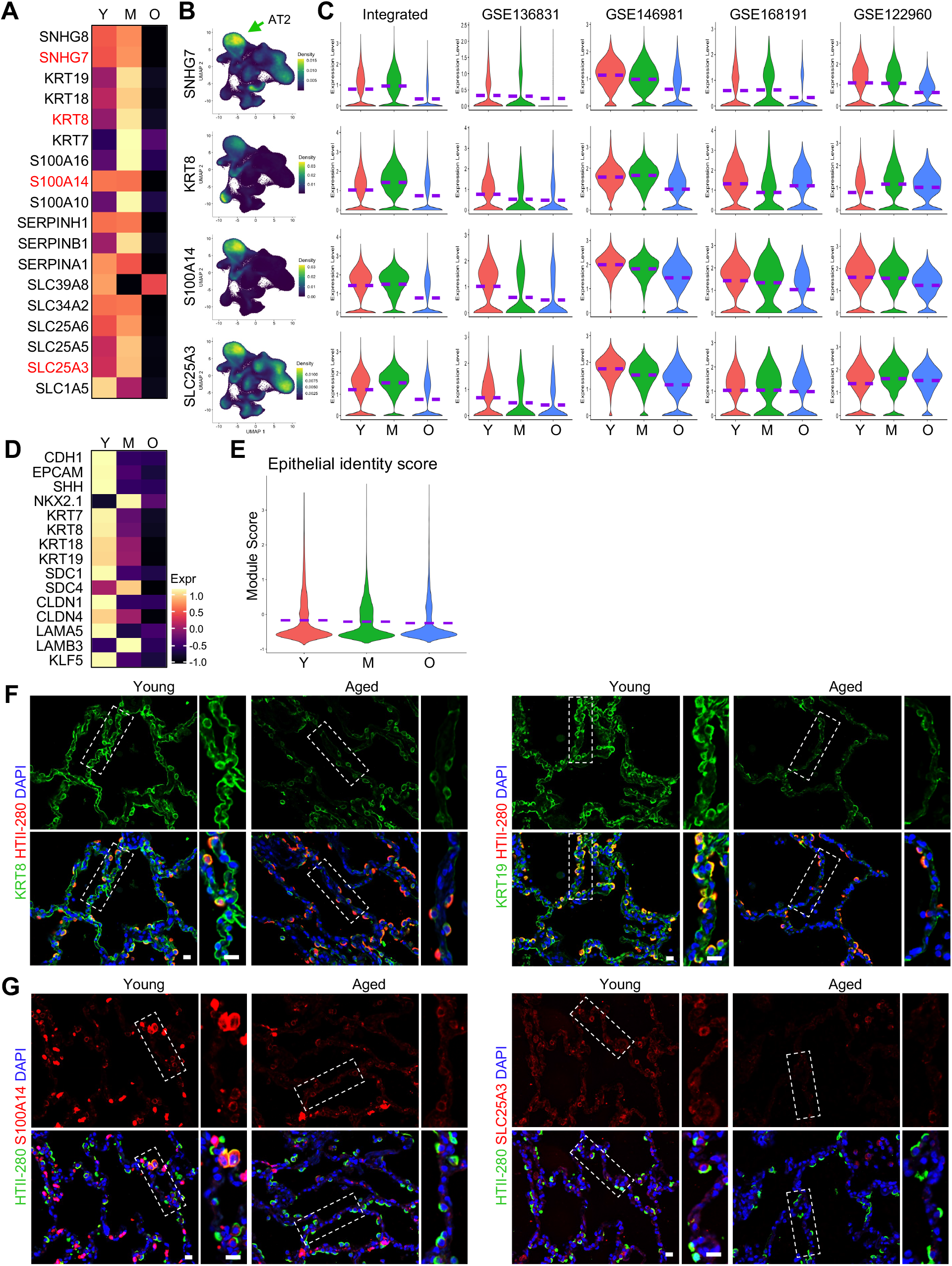
Declined epithelial features in aged human AT2 cells. Related to Figure 2. **(A)** Heatmaps presented the expression of the representative genes downregulated in aged AT2 cells. **(B)** Density plot visualization of the expression and specification of the representative genes (red highlighted in **A**) downregulated in aged AT2 cells. Green arrows indicated the AT2 cell clusters. **(C)** Violin plot visualization of transcriptions of the genes downregulated in aged AT2s in the integrated and representative individual datasets. (**D** and **E**) Heatmaps compared the relative transcriptions of the epithelial identity genes (**D**), and violin plot showed the average expression of the epithelial identity genes (**E**) in AT2 from subjects of different age stages. **(F** and **G)** Immunofluorescence co-staining of epithelial identity gene, KRT8/KRT19 (**F**), and representative aged AT2 downregulated genes, S100A14/SLC25A3 (**G**), with human AT2 cell marker (HTII-280) in alveolar regions of human lung sections from young and aged donors. Boxed regions were magnified. Purple dashed lines in the violin plots indicated the mean levels of gene transcriptions. Y, young donors (Age<40), M, middle-aged donors (Age40-60), O, aged (old) donors (Age>60). Scale bar: 20 µm (**F**, **G**).

**Figure S5.**
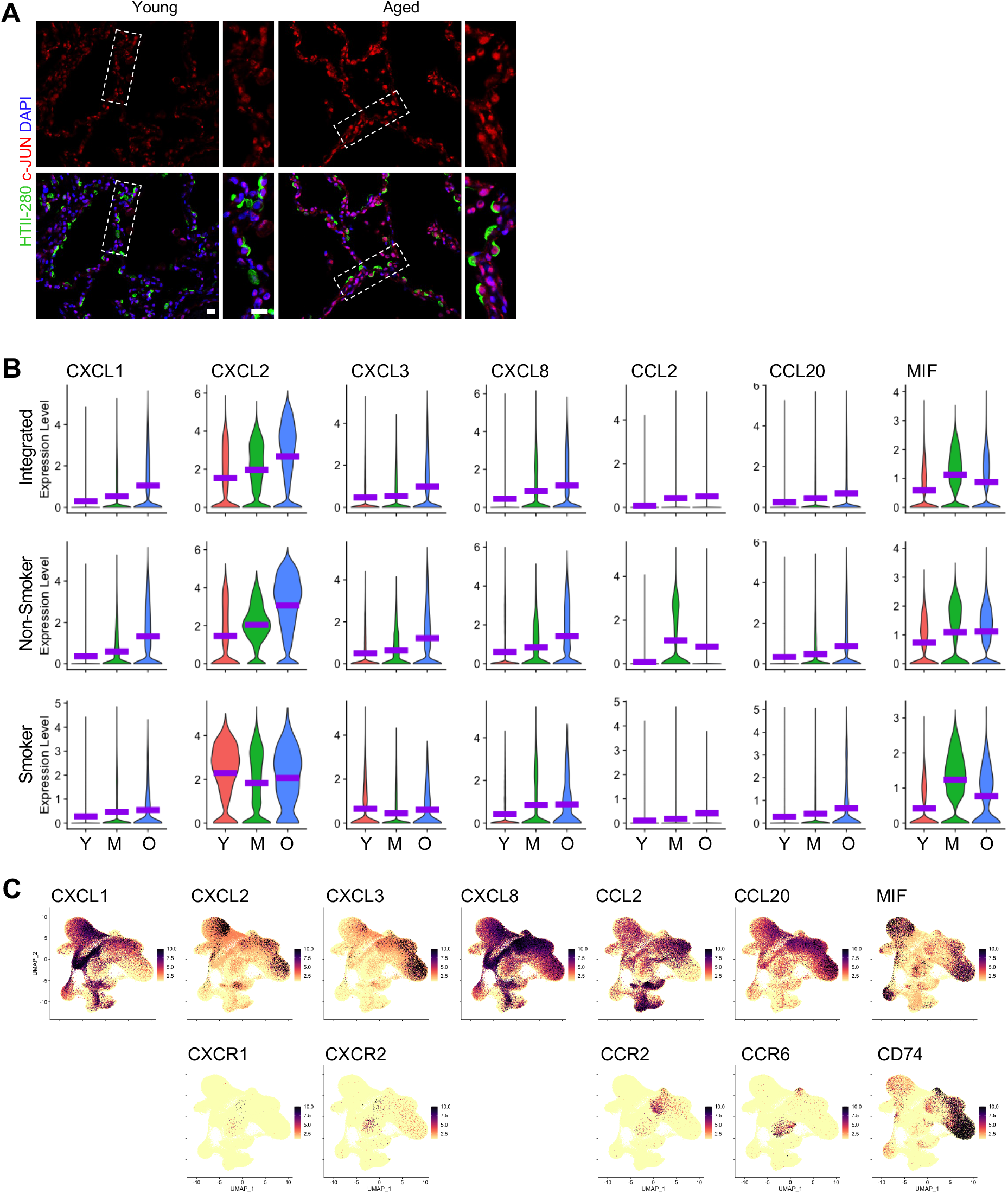
Elevated inflammation in aged human AT2 cells. Related to Figure 2. **(A)** Immunofluorescence staining presented the protein levels of representative AP-1 transcription factor gene, c-JUN/JUN, in human AT2 cells (HTII-280+) on human lung sections from young and aged donors. Boxed regions were magnified. **(B)** Violin plots showed the transcriptions of chemokine genes in AT2 cells from subjects of different age stages from the integrated data, isolated non-smoker dataset, and smoker dataset. All five chemokines showed consistently elevated transcriptions in aged AT2 cells from the integrated data, isolated non-smoker dataset, and smoker dataset. **(C)** UMAP visualization of the transcriptions of the chemokine/cytokine genes and their matched receptor genes. Most of the receptors had limited expression. Purple dashed lines in the violin plots indicated the mean levels of gene transcriptions. Y, young donors (Age<40), M, middle-aged donors (Age40-60), O, aged (old) donors (Age>60). Scale bar: 20 µm (**A**).

**Figure S6.**
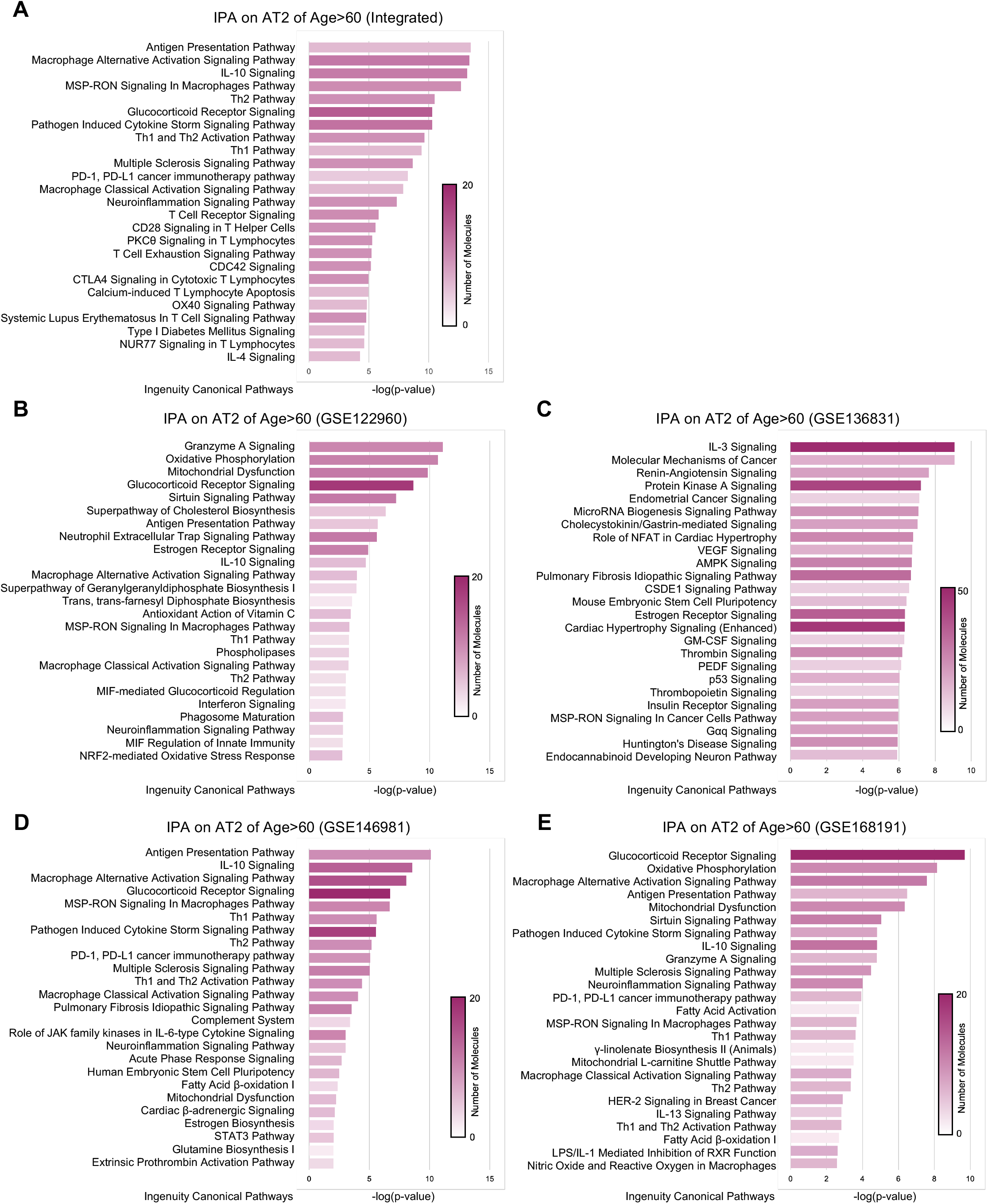
Ingenuity Pathway Analysis (IPA) revealed activated inflammatory pathways in the aged AT2 cells. Related to Figure 2. **(A-E)** Ingenuity Pathway Analysis (IPA) on the aged AT2 upregulated genes from the integrated dataset (**A**) and representative individual datasets (**B-E**) revealed several activated inflammatory pathways in the aged human AT2 cells. The top 25 canonical pathways of each analysis were illustrated. The pathways were ranked by the -log(p-value) and the color scales indicated the number of target molecules (genes) involved in indicated canonical pathways.

**Figure S7.**
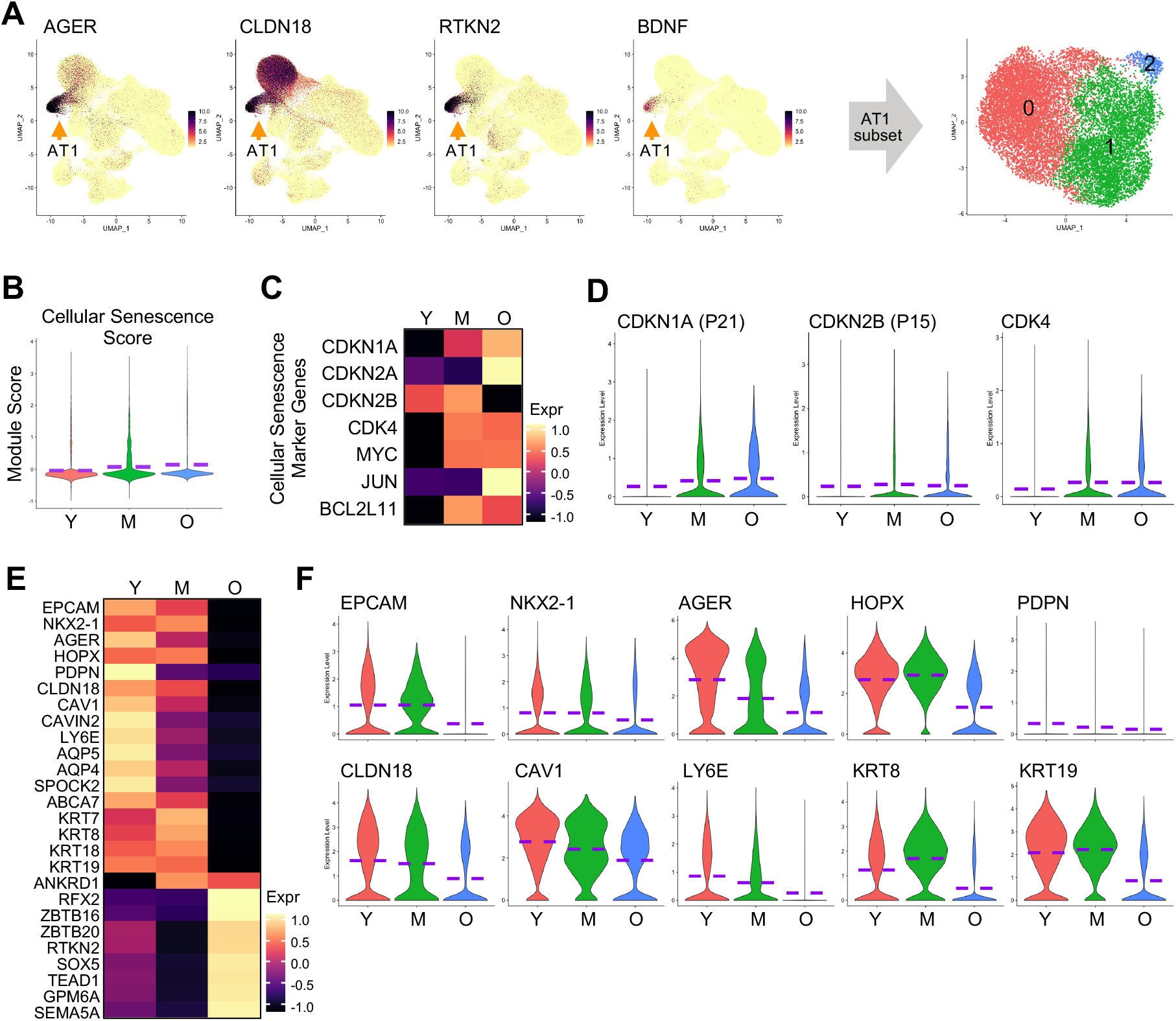
Dysfunctions of AT1 cells from aged subjects. Related to Figure 2. **(A)** AT1 cell cluster was identified in the integrated dataset by checking the expression of canonical AT1 marker genes and was isolated and re-clustered. **(B)** Violin plots presented the cellular senescence scores of AT1 cells from different age stages. The cellular senescence score was defined as the average transcription of 149 core senescence genes from CSGene that were detectable in the AT1 cells. **(C)** Heatmaps showed the transcriptions of representative cellular senescence genes in AT1 cells from different age stages. **(D)** The relative expression levels of representative cellular senescence genes in AT1 cells from subjects of different age stages were visualized by violin plots. **(E)** Heatmaps showed dysregulated gene expression programs of AT1 cells from aged subjects. **(F)** Violin plots showed the transcriptional levels of representative epithelial cell signature genes and AT1 cell functional genes in AT1 cells from subjects of different age stages. Purple dashed lines in the violin plots indicated the mean levels of module scores or gene transcriptions. Orange arrows indicated the AT1 cell clusters (**A**). Y, young donors (Age<40), M, middle-aged donors (Age40-60), O, aged (old) donors (Age>60).

**Figure S8.**
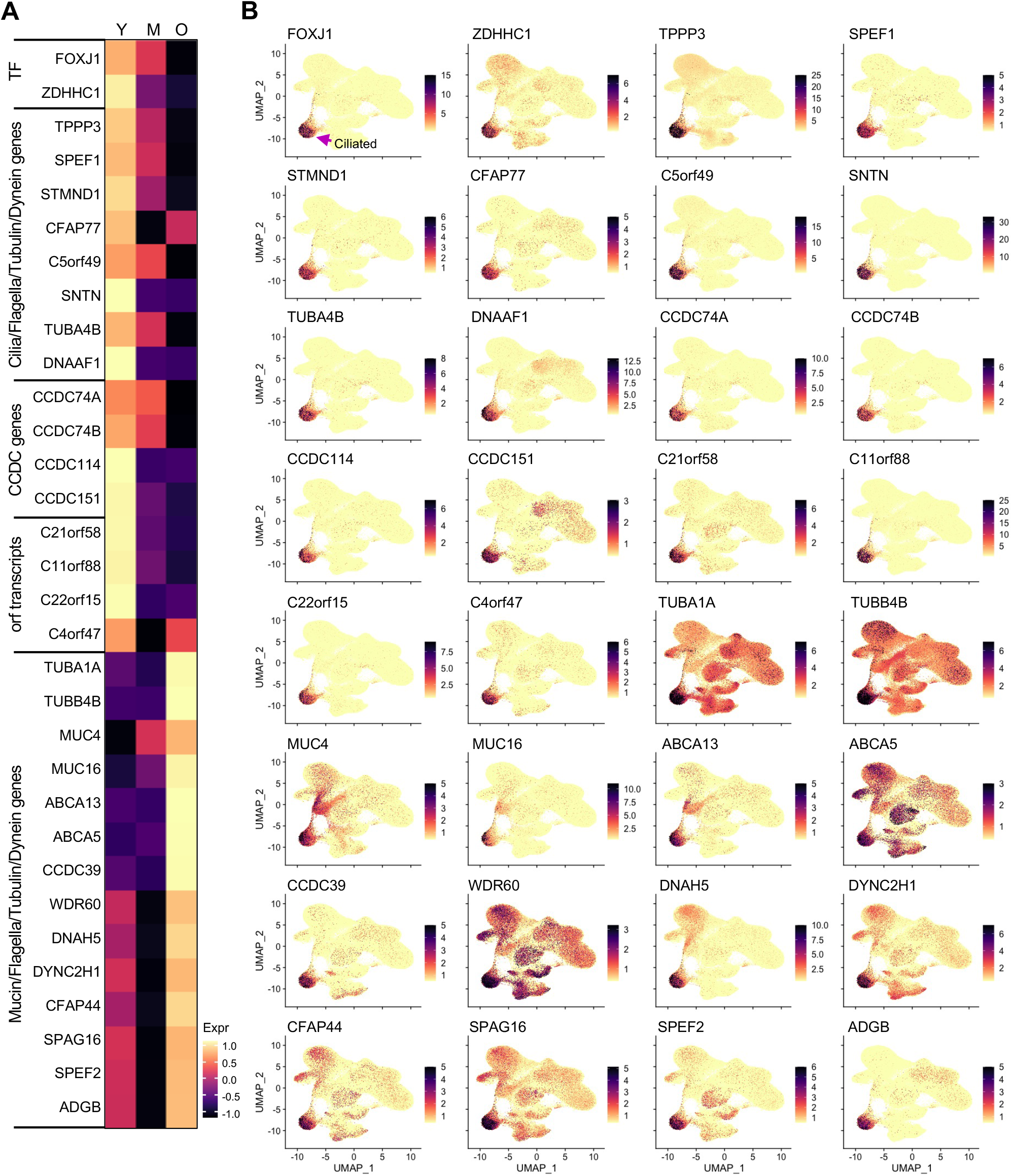
Dysregulated gene expression patterns in ciliated cells from aged subjects. Related to Figure 2. **(A)** Heatmaps presented the relative transcriptions of the ciliated cell specific transcriptional factor genes, known and unknown functional genes in isolated ciliated cells from subjects of different age stages. **(B)** UMAPs showed the specifications of the genes that were dysregulated in ciliated cells from aged subjects. Purple arrow indicated the ciliated cell cluster. TF, transcriptional factor. Y, young donors (Age<40), M, middle-aged donors (Age40-60), O, aged (old) donors (Age>60).

**Figure S9.**
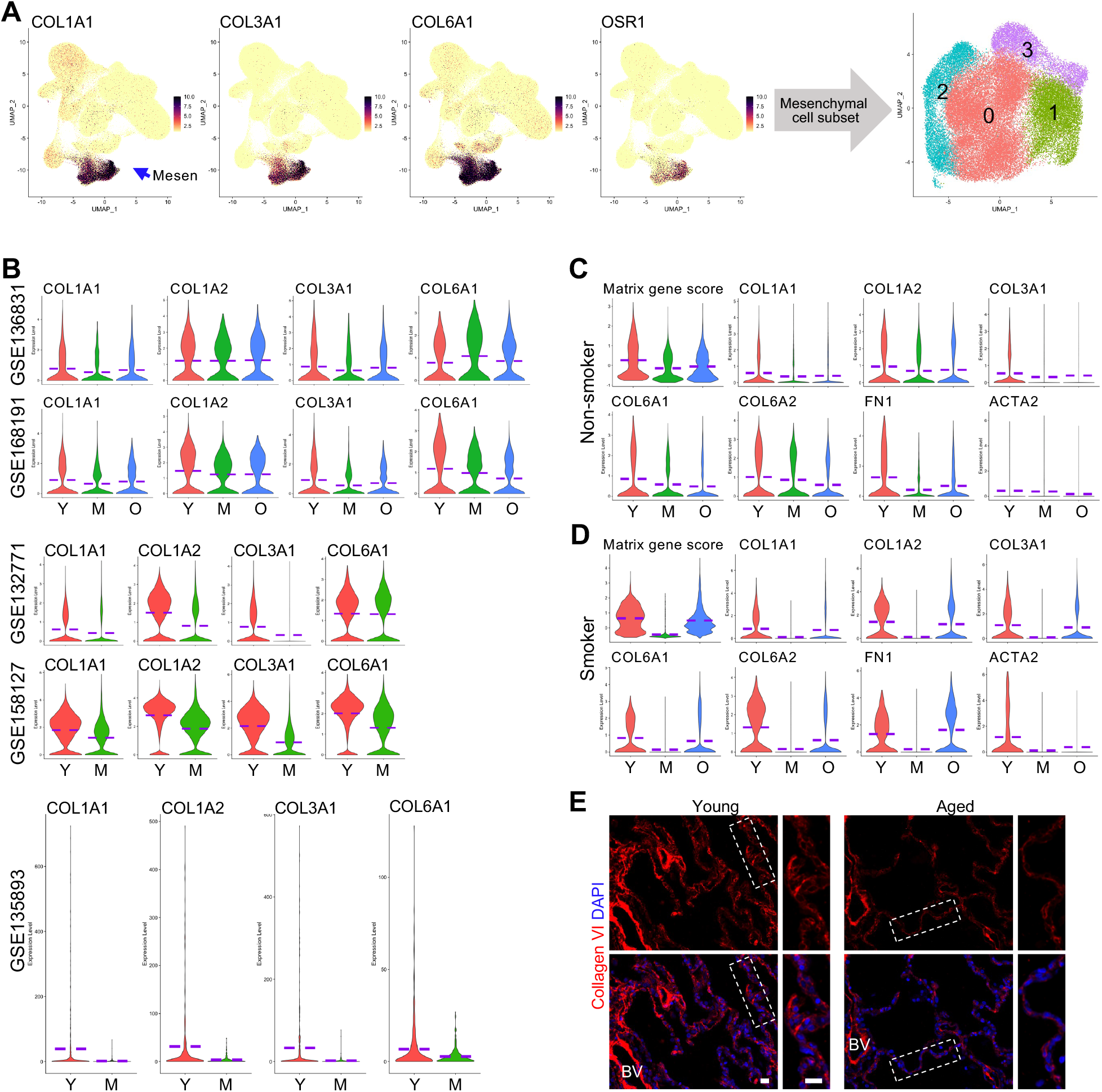
Decreased matrix gene transcription in lung mesenchymal cells of aged subjects. Related to Figure 3. **(A)** Identification and subset of lung mesenchymal cells by checking the expression of canonical mesenchymal marker genes. Blue arrow indicated the mesenchymal cell clusters with specific expression of these genes in the integrated dataset. **(B)** The transcriptions of representative matrix genes in mesenchymal cells from subjects of different age stages in individual datasets were visualized by violin plots. **(C** and **D)** Violin plots presented the matrix gene scores and collagen gene transcriptions in the isolated mesenchymal cells from non-smoking (**C**) and smoking (**D**) donors of different age stages. **(E)** Immunofluorescence staining for Collagen VI on human lung sections from young and aged donors. Boxed regions were magnified. BV, blood vessel. Scale bar: 20 µm. Purple dashed lines in the violin plots indicated the mean levels of module scores and gene transcriptions. Matrix gene score, average expression of COL1A1, COL1A2, COL3A1, COL6A1, COL6A2, FN1 and ACTA2. Y, young donors (Age<40), M, middle-aged donors (Age40-60), O, aged (old) donors (Age>60).

**Figure S10.**
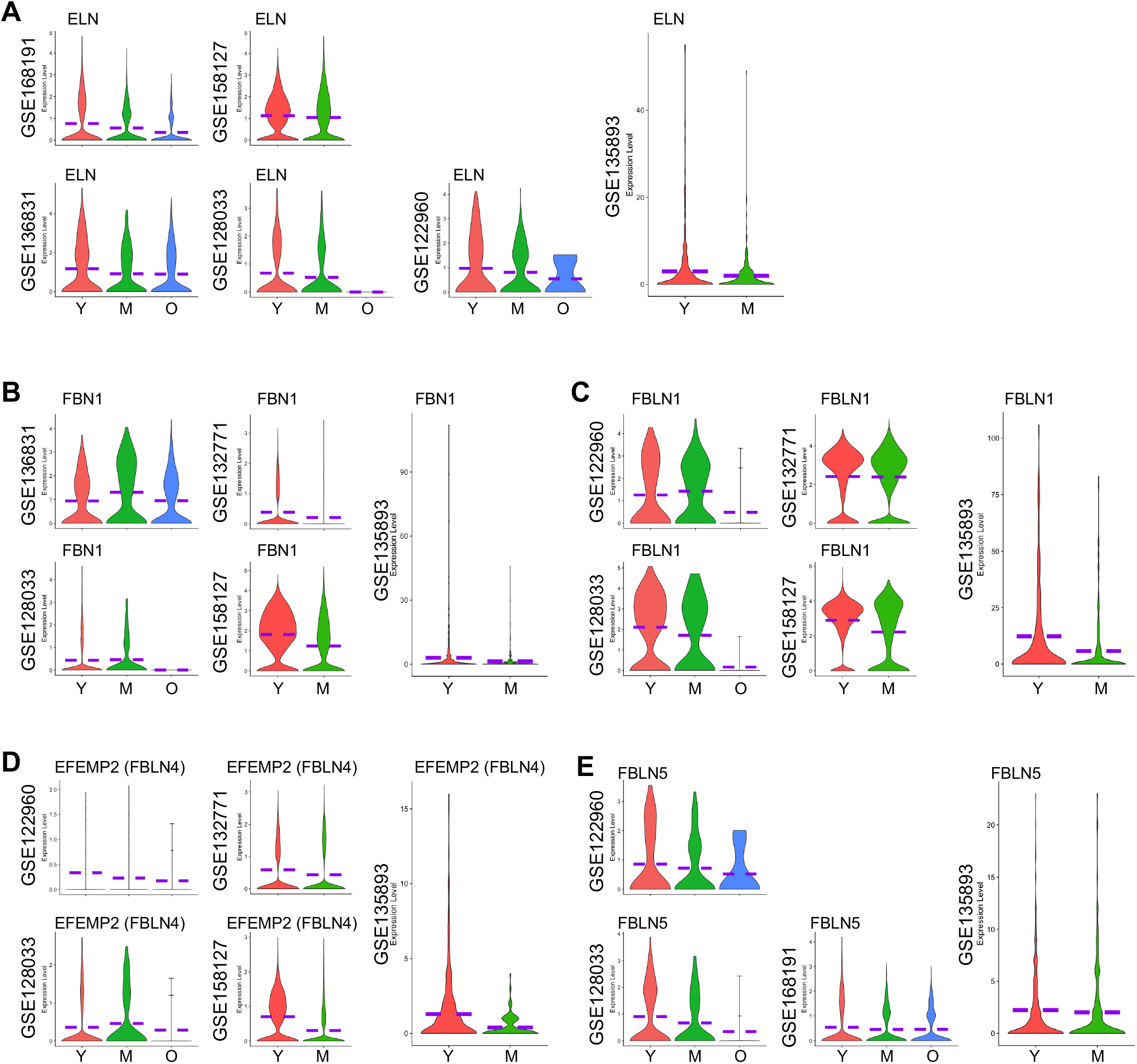
Relative transcriptions of genes specific to elastic fiber components and assembly in aged lung mesenchymal cells in individual datasets. Related to Figure 3. Violin plot showed the transcriptions of genes specific to elastic fiber components and assembly, ELN (**A**), FBN1 (**B**), FBLN1 (**C**), EFEMP2 (FBLN4) (**D**), and FBLN5 (**E**), in mesenchymal cells from subjects of different age stages in different individual datasets. Purple dashed lines in the violin plots indicated the mean levels of gene transcriptions. Y, young donors (Age<40), M, middle-aged donors (Age40-60), O, aged (old) donors (Age>60).

**Figure S11.**
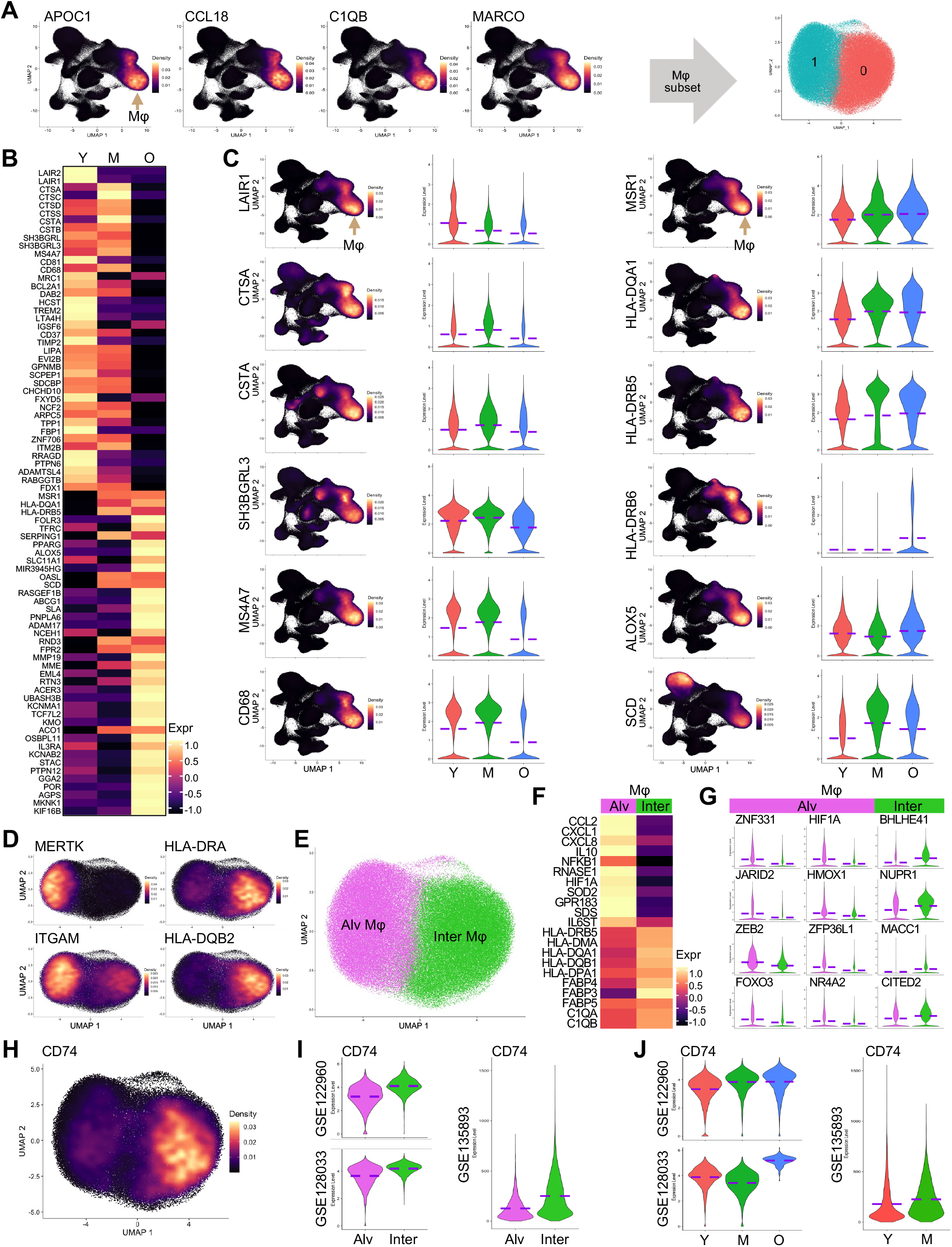
Differential expression analysis of lung macrophages from subjects of different age stages. Related to Figure 4. **(A)** UMAPs showed the transcriptions of macrophage marker genes and the isolation and re-clustering of macrophages. **(B)** Visualization of the top 40 genes dysregulated in young and aged lung macrophages by heatmaps. **(C)** Density and violin plots revealed the specification and dysregulation of functional genes specific to macrophages in lungs from aged subjects. **(D)** Density plots showed the transcriptions of alveolar and interstitial macrophage specific genes in the clusters of isolated lung macrophages. **(E)** Alveolar and interstitial macrophage clusters were identified based on the gene transcriptions in (**D**). **(F** and **G)** Heatmap and violin plot visualization of relative transcriptional levels of the differentially expressed genes (**F**) and transcription factors (**G**) specific to alveolar and interstitial macrophages, respectively. **(H)** Density plot presented the expression of MIF receptor gene, CD74, in macrophage clusters. **(I** and **J)** Violin plots showed the relative expression of CD74 in alveolar vs interstitial Mφ (**I**) and in total lung Mφ (**J**) from different age stages. Y, young donors (Age<40), M, middle-aged donors (Age40-60), O, aged (old) donors (Age>60). Mφ, macrophages. Brown arrow indicated macrophage (Mφ) clusters.

**Figure S12.**
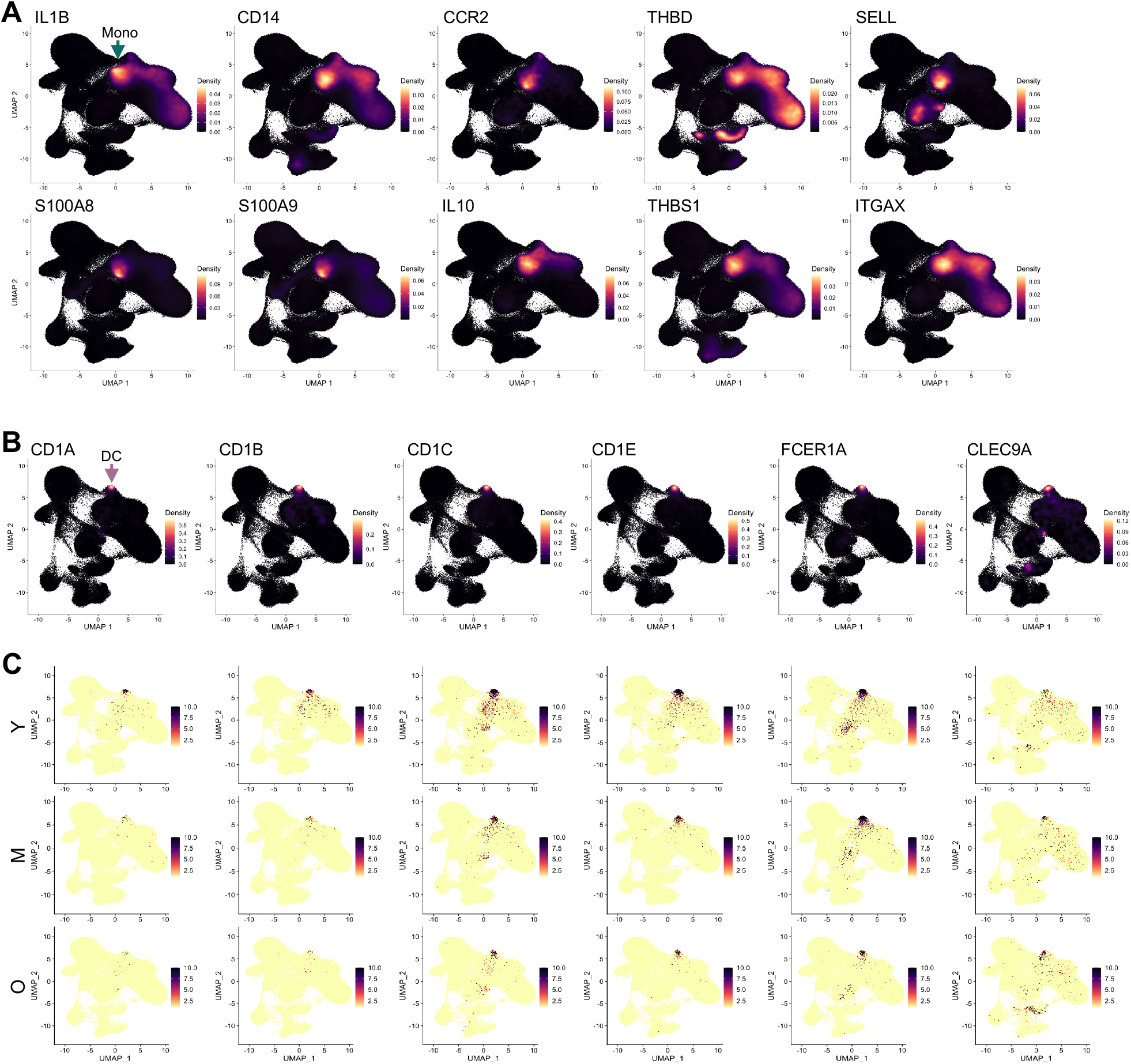
Differential expression analysis of lung monocytes and DCs from subjects of different age stages. Related to Figure 4. **(A)** Density plots showed the specification of cytokine genes in lung monocytes. Teal arrow indicated monocyte (Mono) clusters. **(B)** Density plots visualized in UMAPs indicated the specification of cell surface antigen/receptor genes in lung DCs. Mauve arrow indicated dendritic cell (DC) cluster. **(C)** Split UMAPs compared the transcription levels of the above genes (**B**) in DCs from subjects of different age stages. Y, young donors (Age<40), M, middle-aged donors (Age40-60), O, aged (old) donors (Age>60).

**Figure S13.**
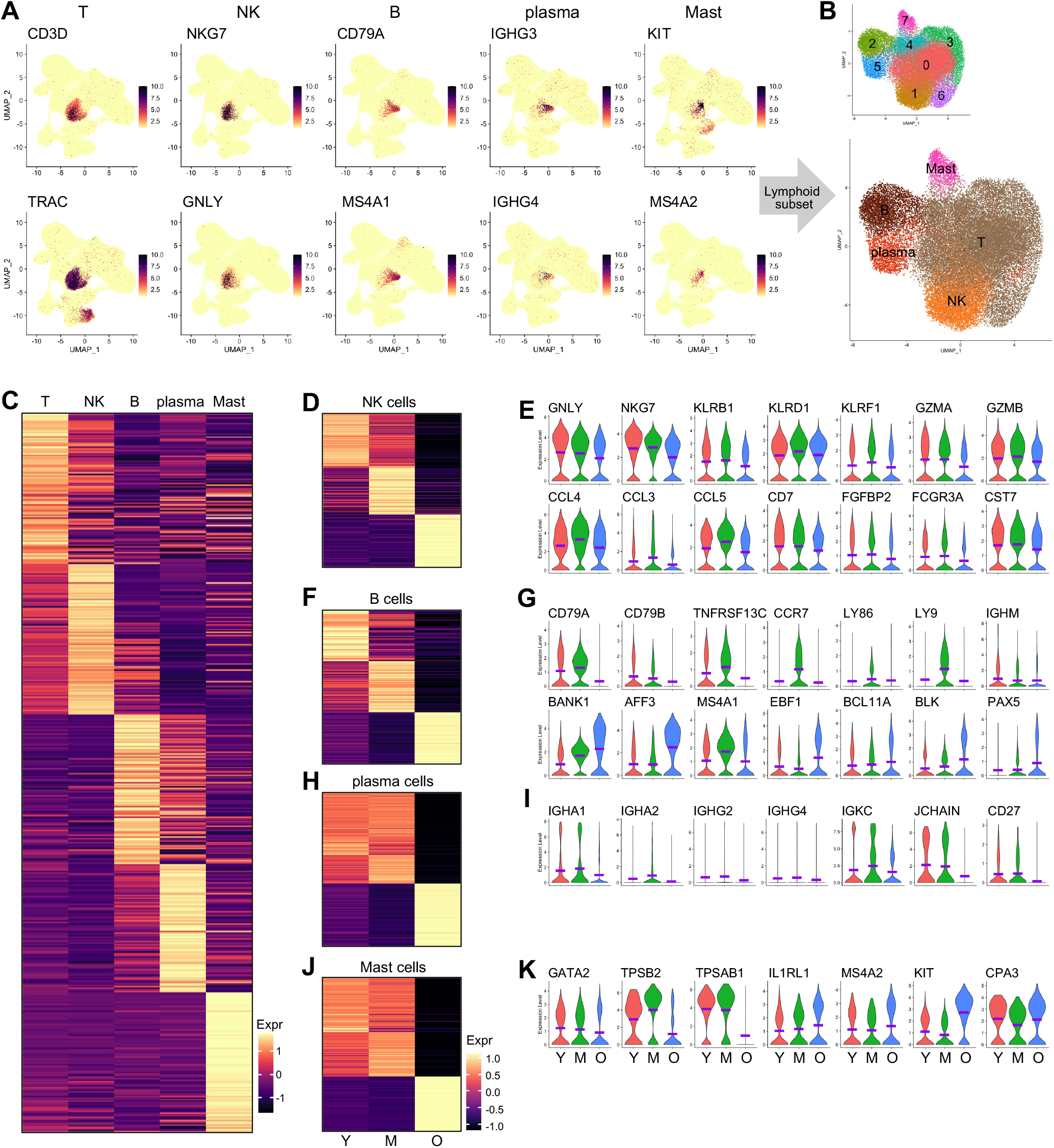
Differential analyses of lymphoid lineages from subjects of different age stages. Related to Figure 4. **(A-B)** Subsets of lymphoid cell lineages and mast cells were identified by checking the expression of canonical marker genes (**A**) and were isolated, re-clustered, and annotated from the integrated data (**B**). **(C)** Heatmap showed the transcriptions of top 100 genes (rows, ranked by log fold-change of the average expression) of each cell type (columns). **(D**, **F**, **H**, and **J)** Differential analysis revealed the top 100 differentially expressed genes (rows, ranked by log fold-change of the average expression) in NK (**D**), B (**F**), Plasma (**H**), and Mast (**J**) cells from lungs of different age stages. Heatmaps shared the same color key. **(E**, **G**, **I**, and **K)** Violin plots showed the dysregulation of canonical functional/differentiation genes in NK (**E**), B (**G**), Plasma (**I**), and Mast (**K**) cells from lungs of aged subjects. In the NK (**E**) and Plasma (**I**) cells all the listed genes were downregulated in aged subjects, while in B (**G**), and Mast (**K**) cells both representative downregulated and upregulated genes were illustrated. Purple dashed lines in the violin plots indicated the mean levels of gene transcriptions. Y, young donors (Age<40), M, middle-aged donors (Age40-60), O, aged (old) donors (Age>60).

**Figure S14.**
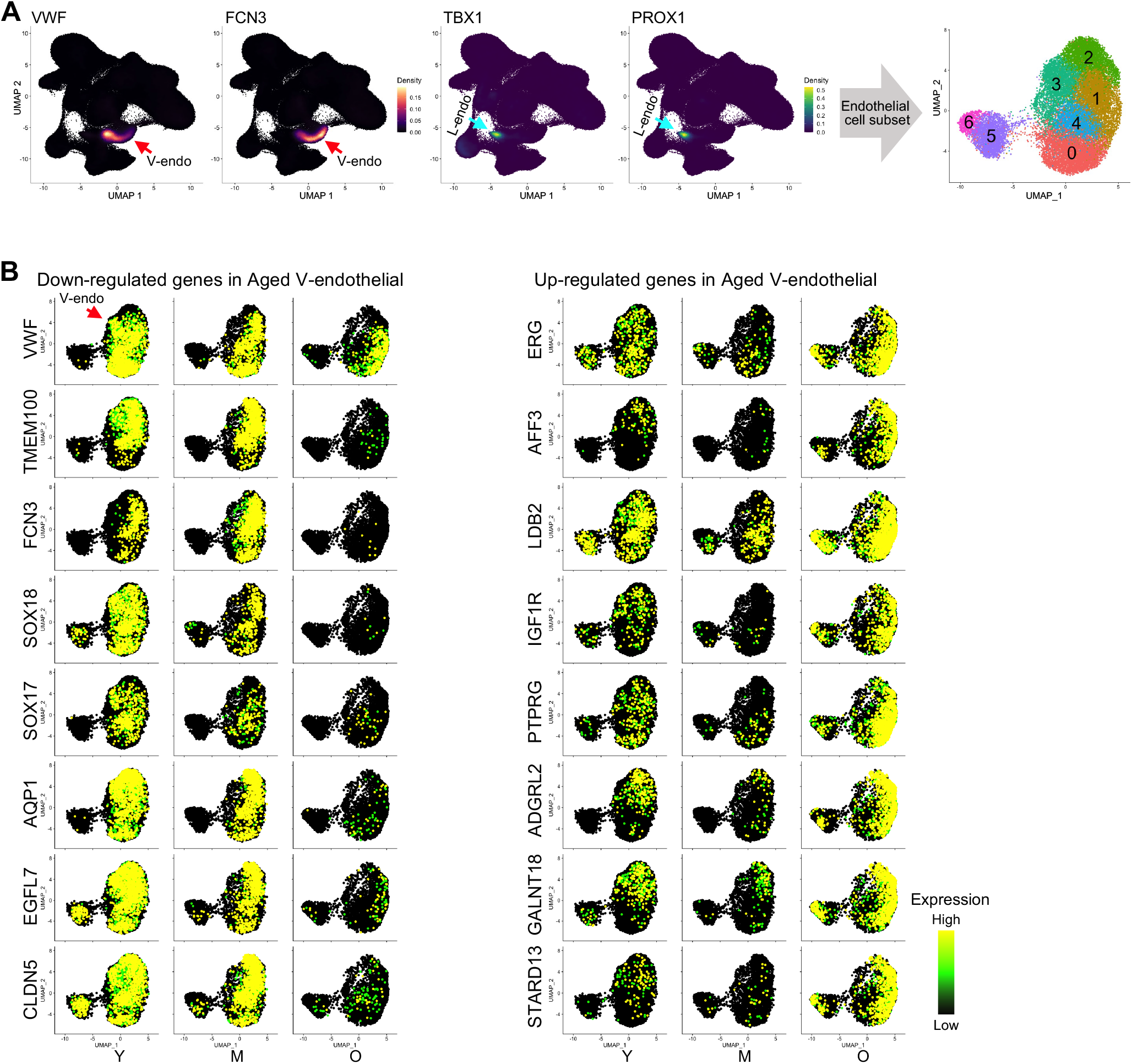
Dysfunctions of vascular endothelial cells from lungs of aged subjects. Related to Figure 5. **(A)** Lung vascular and lymphatic endothelial cell clusters were identified by checking the expression of canonical marker genes and were isolated from the integrated data. Red and cyan arrows indicated the V- and L-endothelial clusters, respectively. **(B)** UMAP visualization of the downregulated functional genes (left panels) and upregulated genes (right panels) in vascular endothelial cells from aged donors. Red arrow indicated the V-endothelial cluster. V-endo, vascular endothelial cells, L-endo, lymphatic endothelial cells. Y, young donors (Age<40), M, middle-aged donors (Age40-60), O, aged (old) donors (Age>60).

**Figure S15.**
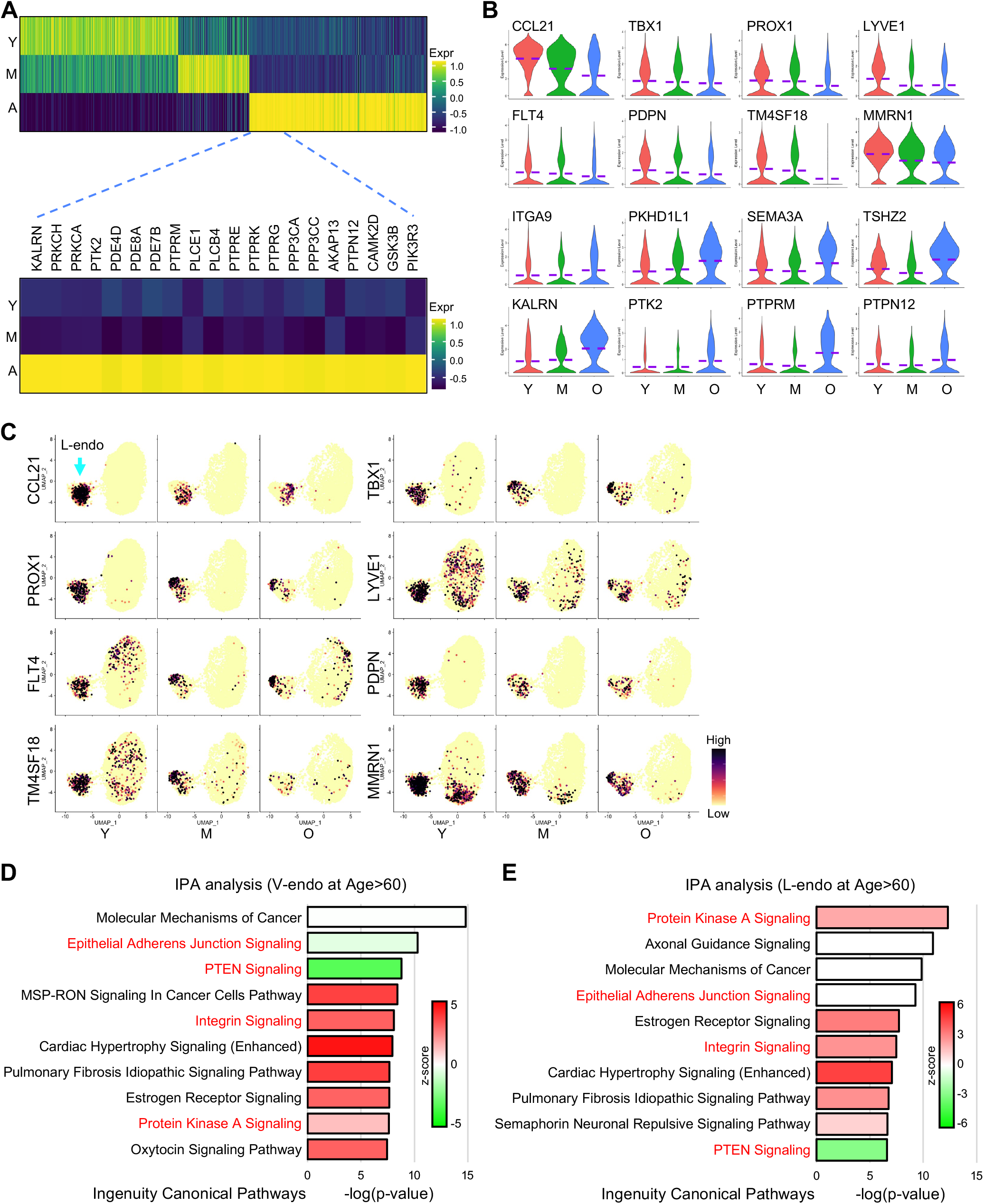
Hypofunctions of lymphatic endothelial cells in aged human lungs. Related to Figure 5. **(A)** Heatmap visualization of the top differentially expressed genes (columns, ranked by log fold-change of the average expression) in the lymphatic endothelial cells from donors at different age stages (rows). Top upregulated genes in aged subjects were listed. **(B)** Transcriptions of the representative dysregulated functional genes in lymphatic endothelial cells from lungs of aged subjects. **(C)** UMAPs showed the transcriptions of the functional genes in lymphatic endothelial cells from subjects of different age stages. Cyan arrow indicated the L-endothelial cluster. **(D** and **E)** Top canonical pathways of genes upregulated in aged vascular (**D**) and lymphatic (**E**) endothelial cells identified by Ingenuity Pathway Analysis (IPA) Purple dashed lines in the violin plots indicated the mean levels of gene transcriptions. L-endothelial, lymphatic endothelial cells. Y, young donors (Age<40), M, middle-aged donors (Age40-60), O, aged (old) donors (Age>60).

**Figure S16.**
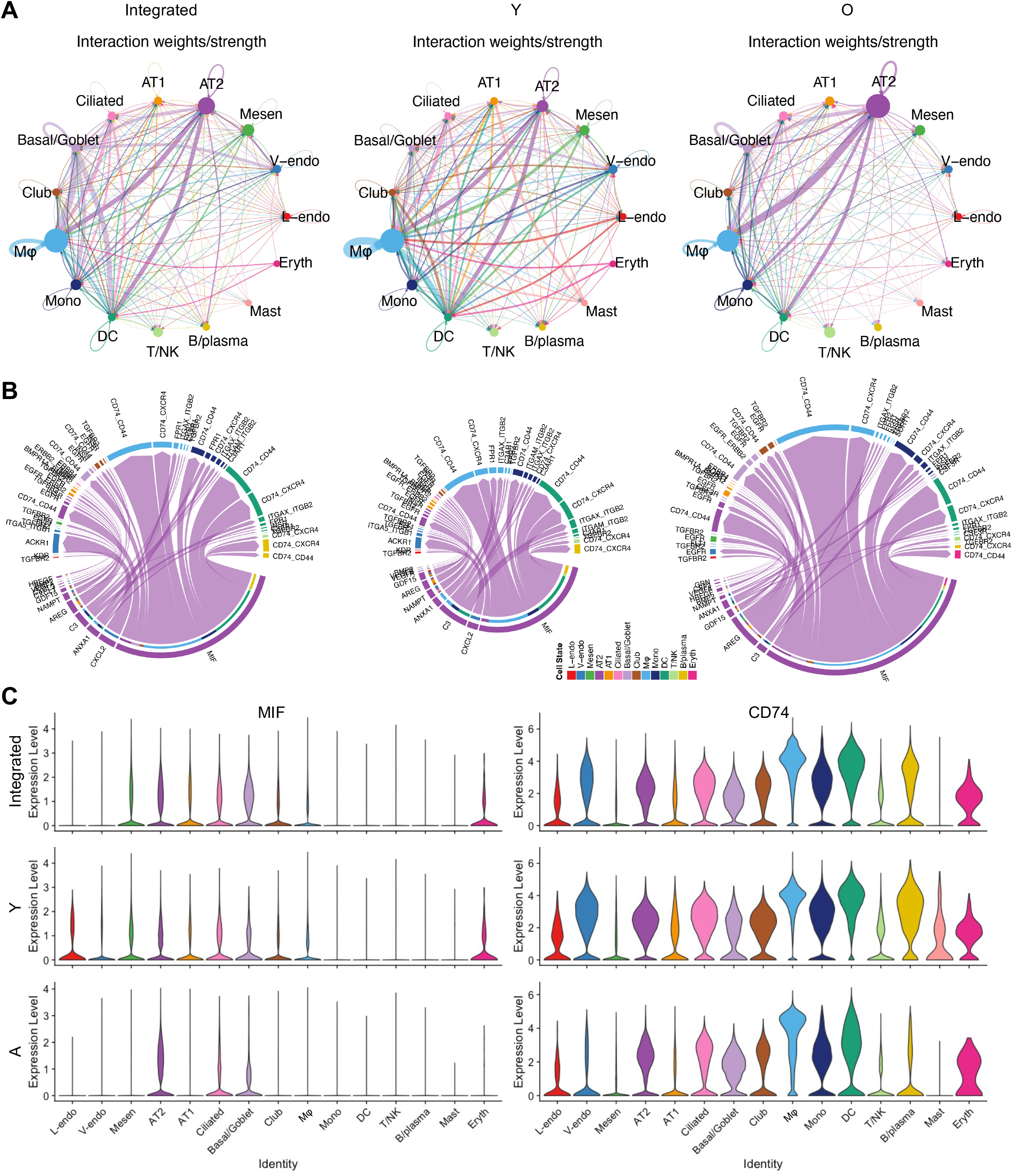
Inference of the cell-cell communications at a signaling pathway level by CellChat. Related to Figure 6. **(A)** Circle plot visualization of the aggregated cell-cell communication networks showed the total interaction strength (weights) among different cell types in the integrated, young (Y) and aged (O) datasets. **(B)** Chord diagrams showed all the significant interactions (L-R pairs) from AT2 to the other cell types in the integrated, young (Y) and aged (O) datasets. The diameters of the Chord diagrams indicated the sum strength (weights) of the interactions. **(C)** Violin plots presented the transcription levels of the top significant interaction L-R pair genes, MIF-CD74, in the cell types of the integrated, young (Y) and aged (O) datasets. L/V-endo, lymphatic/vascular endothelial cells, Mesen, mesenchymal cells, AT1/2, alveolar type I/II epithelial cells, Mφ, macrophages, Mono, monocytes, DC, dendritic cells, Eryth, erythrocytes. Y, young donors (Age<40), M, middle-aged donors (Age40-60), O, aged (old) donors (Age>60).

**Table S1.**
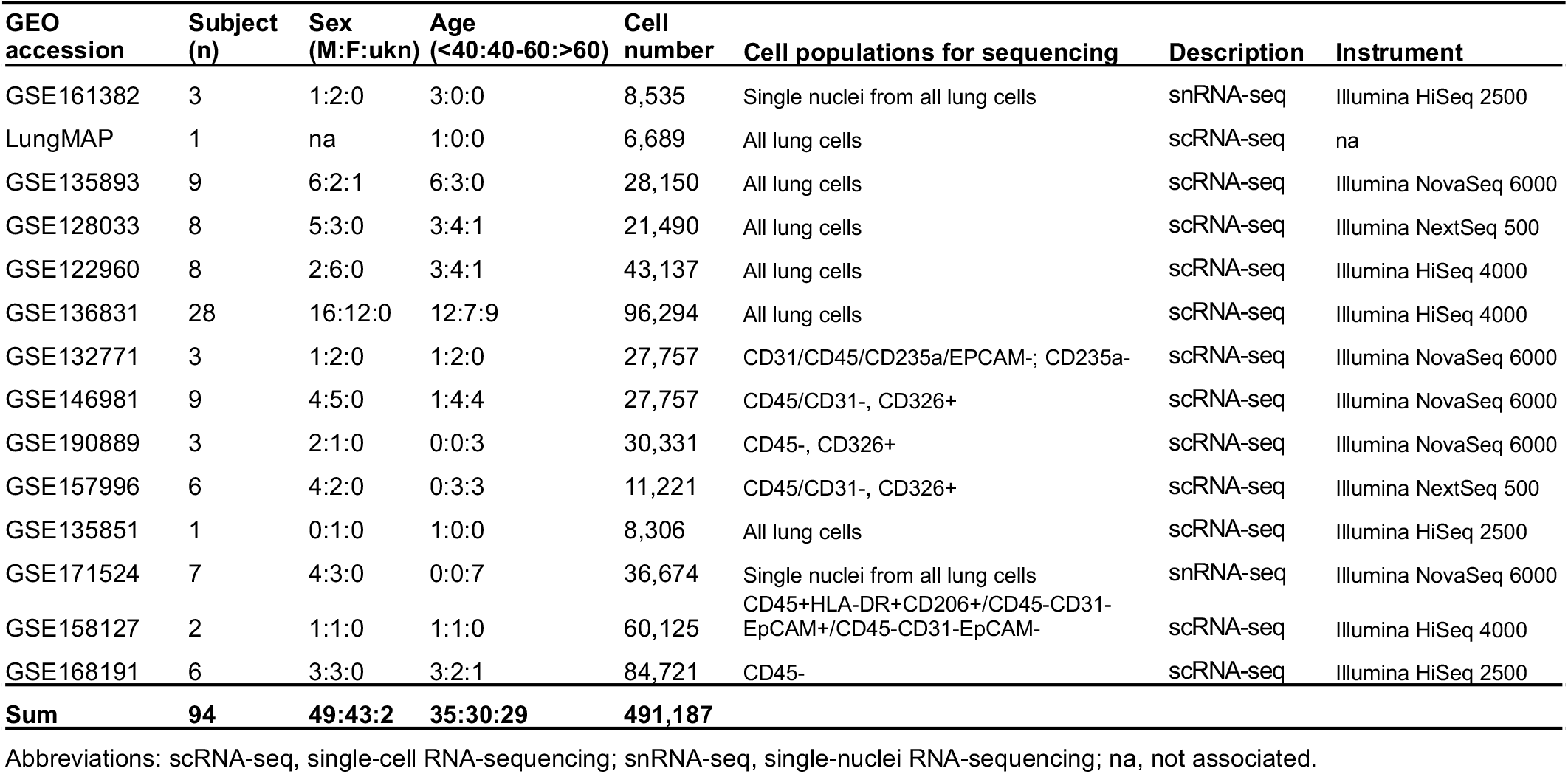
Details of the scRNA/snRNA-seq datasets used for the integration.

| **GEO accession** | **Subject (n)** | **Sex (M:F:ukn)** | **Age (<40:40-60:>60)** | **Cell number** | **Cell populations for sequencing** | **Description** | **Instrument** |
| --- | --- | --- | --- | --- | --- | --- | --- |
| GSE161382 | 3 | 1:2:0 | 3:0:0 | 8,535 | Single nuclei from all lung cells | snRNA-seq | Illumina HiSeq 2500 |
| LungMAP | 1 | na | 1:0:0 | 6,689 | All lung cells | scRNA-seq | na |
| GSE135893 | 9 | 6:2:1 | 6:3:0 | 28,150 | All lung cells | scRNA-seq | Illumina NovaSeq 6000 |
| GSE128033 | 8 | 5:3:0 | 3:4:1 | 21,490 | All lung cells | scRNA-seq | Illumina NextSeq 500 |
| GSE122960 | 8 | 2:6:0 | 3:4:1 | 43,137 | All lung cells | scRNA-seq | Illumina HiSeq 4000 |
| GSE136831 | 28 | 16:12:0 | 12:7:9 | 96,294 | All lung cells | scRNA-seq | Illumina HiSeq 4000 |
| GSE132771 | 3 | 1:2:0 | 1:2:0 | 27,757 | CD31/CD45/CD235a/EPCAM-; CD235a- | scRNA-seq | Illumina NovaSeq 6000 |
| GSE146981 | 9 | 4:5:0 | 1:4:4 | 27,757 | CD45/CD31-, CD326+ | scRNA-seq | Illumina NovaSeq 6000 |
| GSE190889 | 3 | 2:1:0 | 0:0:3 | 30,331 | CD45-, CD326+ | scRNA-seq | Illumina NovaSeq 6000 |
| GSE157996 | 6 | 4:2:0 | 0:3:3 | 11,221 | CD45/CD31-, CD326+ | scRNA-seq | Illumina NextSeq 500 |
| GSE135851 | 1 | 0:1:0 | 1:0:0 | 8,306 | All lung cells | scRNA-seq | Illumina HiSeq 2500 |
| GSE171524 | 7 | 4:3:0 | 0:0:7 | 36,674 | Single nuclei from all lung cells | snRNA-seq | Illumina NovaSeq 6000 |
| GSE158127 | 2 | 1:1:0 | 1:1:0 | 60,125 | CD45+HLA-DR+CD206+/CD45-CD31-EpCAM+/CD45-CD31-EpCAM- | scRNA-seq | Illumina HiSeq 4000 |
| GSE168191 | 6 | 3:3:0 | 3:2:1 | 84,721 | CD45- | scRNA-seq | Illumina HiSeq 2500 |
| **Sum** | **94** | **49:43:2** | **35:30:29** | **491,187** |  |  |  |

Abbreviations: scRNA-seq, single-cell RNA-sequencing; snRNA-seq, single-nuclei RNA-sequencing; na, not associated.

